# Delivery determinants of an *Acinetobacter baumannii* type VI secretion system bifunctional peptidoglycan hydrolase

**DOI:** 10.1101/2024.06.29.601107

**Authors:** Valeriya Bezkorovayna, Brooke K. Hayes, Francesca N. Gillett, Amy Wright, David I. Roper, Marina Harper, Sheena McGowan, John D. Boyce

## Abstract

*Acinetobacter baumannii* is a Gram-negative opportunistic pathogen that is a common cause of nosocomial infections. The increasing development of antibiotic resistance in this organism is a global health concern. The *A. baumannii* clinical isolate AB307-0294 produces a type VI secretion system (T6SS) that delivers three antibacterial cargo effector proteins (Tse15, Tde16 and Tae17) that give this strain a competitive advantage against other bacteria in polymicrobial environments. These effectors are delivered via specific non-covalent interactions with the T6SS needle tip proteins VgrG15, VgrG16 and VgrG17, respectively. Here we determine the molecular function of the Tae17 effector protein and define the regions of Tae17 and VgrG17 essential for its delivery. Specifically, we show that Tae17 is a multidomain, bifunctional peptidoglycan-degrading enzyme. Tae17 has both lytic transglycosylase activity, which targets the peptidoglycan sugar backbone, and amidase activity, which targets the sugar-peptide bonds. Moreover, we show that the transglycosylase activity was more important for killing *Escherichia coli*. Using deletion constructs and bacterial two-hybrid analyses, we identify that amino acids 1051-1085 of the VgrG17 needle tip protein and amino acids 1-162 of the Tae17 effector protein are necessary for the Tae17:VgrG17 interaction. Furthermore, we identify the VgrG17 amino acids G1069 and W1075 as crucial for the delivery of Tae17; the first time such specific delivery determinants of T6SS cargo effectors have been defined. This study provides molecular insight into how the T6SS allows *A. baumannii* strains to gain dominance in polymicrobial communities and thus improve their chances of survival and transmission.

**IMPORTANCE:** We have shown that the *Acinetobacter baumannii* T6SS effector Tae17 is a modular, bifunctional, peptidoglycan-degrading enzyme that has both lytic transglycosylase and amidase activity. Both activities contribute to the ability to degrade peptidoglycan, but the glycosyltransferase activity was more important for the interbacterial killing of *Escherichia coli*. We have defined the specific regions of Tae17 and its cognate delivery protein VgrG17 that are necessary for the non-covalent interactions and, for the first time, identified specific amino acids essential for delivery. This work contributes to our molecular understanding of bacterial competition strategies in polymicrobial environments and may provide a window to the design of new therapeutic approaches for combating infection by *A. baumannii*.

## INTRODUCTION

*Acinetobacter baumannii* is a Gram-negative bacterium that is a common cause of hospital-acquired infections. Its ability to withstand hospital disinfection procedures and its increasing resistance to current clinical antibiotics has led to it being listed as one of three pathogens where the need for research and development of new antibiotics is “critical” (1–3). Many strains of *A. baumannii* express a type VI secretion system (T6SS), which plays a primary role in interbacterial competition and its survival in polymicrobial environments (4–6).

The T6SS is found in many Gram-negative bacterial species, especially α-, β- and γ- proteobacteria (7). This molecular “nanomachine”, which resembles the inverted tail of a bacteriophage (8), is anchored in the cell membrane and delivers effector proteins directly into target cells or the extracellular environment. Some T6SS effectors can target eukaryotic cells and therefore play roles in virulence, but many are antibacterial toxins that mediate interbacterial competition. The T6SS comprises a membrane complex, which anchors the system in the bacterial cell (9), an outer sheath, composed of TssB and TssC, and an inner needle composed of repeating Hcp hexamers (10) topped by a spike consisting of a trimer of VgrG proteins (11) and a single PAAR protein (12). Contraction of the outer sheath propels the inner needle, and any effectors attached to the inner needle, outside of the cell.

T6SS effectors can be classified by their mechanism of delivery. Specialized effectors are those that are translationally fused to T6SS needle proteins while cargo effectors are delivered via non-covalent interactions with these proteins (13). Currently, the specific interactions required for the delivery of cargo effectors are poorly understood. Cargo effectors often interact with the VgrG proteins that form the T6SS needle tip. These VgrG proteins have a highly conserved N-terminal VgrG domain and a variable C-terminal region. The N-terminal VgrG domain is crucial for T6SS spike-formation (11, 14) while the C-terminal region is proposed to interact with the cognate effector (15–17). However, there is currently no information on the precise residues critical for the delivery of these cargo effectors.

Since T6SS-mediated killing is contact-dependent (18), this form of interbacterial competition can become self-limited by the formation of “corpse barriers” that provide steric protection from T6SS-mediated attack (19). To overcome this effect, many bacterial species encode at least one lytic T6SS effector that targets bacterial peptidoglycan (PG), an essential component of the bacterial cell wall. PG hydrolase T6SS effectors usually cleave either the glycan linkages between *N*-acetylmuramic acid (MurNAc) and *N*-acetylglucosamine (GlcNAc) (20), or the cross-linking peptide stems of peptidoglycan (4). Currently, only one effector has been described that has two separate catalytic domains and targets both the glycan strands and the interpeptide bonds (21).

The *A. baumannii* strain AB307-0294 (22) encodes a single, constitutively active T6SS that delivers three cargo effectors, Tse15, Tde16 and Tae17. Each effector can act alone and kill competing bacteria, including *Escherichia coli* and *Acinetobacter baylyi* (6). The effector Tae17 is a predicted peptidoglycan hydrolase that is proposed to be delivered via non-covalent interactions with its cognate VgrG protein, VgrG17. This study aimed to functionally characterize Tae17 and define the specific interactions required for its delivery by the T6SS. Here we show that the C-terminal region of VgrG17 interacts with the N-terminal region of the Tae17 effector and for the first time identify specific amino acids within the C-terminal region of VgrG17 that are essential for this interaction. Furthermore, we show that Tae17 is a multi-domain, bifunctional peptidoglycan-degrading effector containing both lytic transglycosylase and amidase activity that are both required for maximal toxicity.

## RESULTS

### Tae17 is a modular, multi-domain effector with both lytic transglycosylase and amidase activity

Previous bioinformatic analysis of the wild-type Tae17 effector, identified a putative peptidoglycan binding LysM domain between residues 112-155, suggesting that Tae17 acts as a peptidoglycan hydrolase (6, 23). Tae17 showed limited amino acid sequence identity to proteins outside the *Acinetobacter* genus (best hit 35% identity with 62% coverage to a LysM domain-containing protein from *Psychrobacter maritimus*) and no significant shared amino acid sequence identity to any functionally characterized proteins. To predict other potential domains and active site residues, we produced a molecular model of wild-type Tae17 using AlphaFold2. The model predicted with high confidence that Tae17 contained four domains (residues 1–88, 110–156, 234–446 and 469–582) each joined by a poorly modeled linker region (Fig. 1A and B). The N-terminal domain was predicted to form an Ig-like fold (Fig. 1C), which is a domain found in a diverse set of proteins that often facilitates protein-protein interactions (12, 24). The second domain, predicted to be a LysM domain, aligned well with characterized LysM structures (25). Using the DALI server for comparison to known structures (26), the third domain (residues 234-446) shared structural homology with a domain in the lytic transglycosylase Cj0843c from *Campylobacter jejuni* (PDB ID: 7LAM, overall RMSD: 20.8 Å, active site RMSD: 11.0 Å) (27). While the overall alignment between the third domain of Tae17 and 7LAM was poor, likely due to the difference in the length of the proteins, alignment of the active site could be visualized. The key active site residues for Cj0843c are R388, E390 and K505, which aligned closely with Tae17 residues H276, E278 and K436 respectively (Fig. 1D). The fourth domain of Tae17 (residues 469-582) aligned with the amidase domain of the T6SS effector Tae3 from *Ralstonia pickettii* (PDB ID: 4HZ9, RMSD: 2.2 Å) (28). The catalytic dyad of Tae3 is known to include residues C23 and H81, closely aligning with Tae17 residues C488 and H550, respectively (Fig. 1E). These residues are consistent with the position of the active site residues in other proteins belonging to the CHAP amidase protein family (29). Together these data suggested that Tae17 may contain a lytic transglycosylase and an amidase domain, both with distinct active sites.

**FIG 1.**
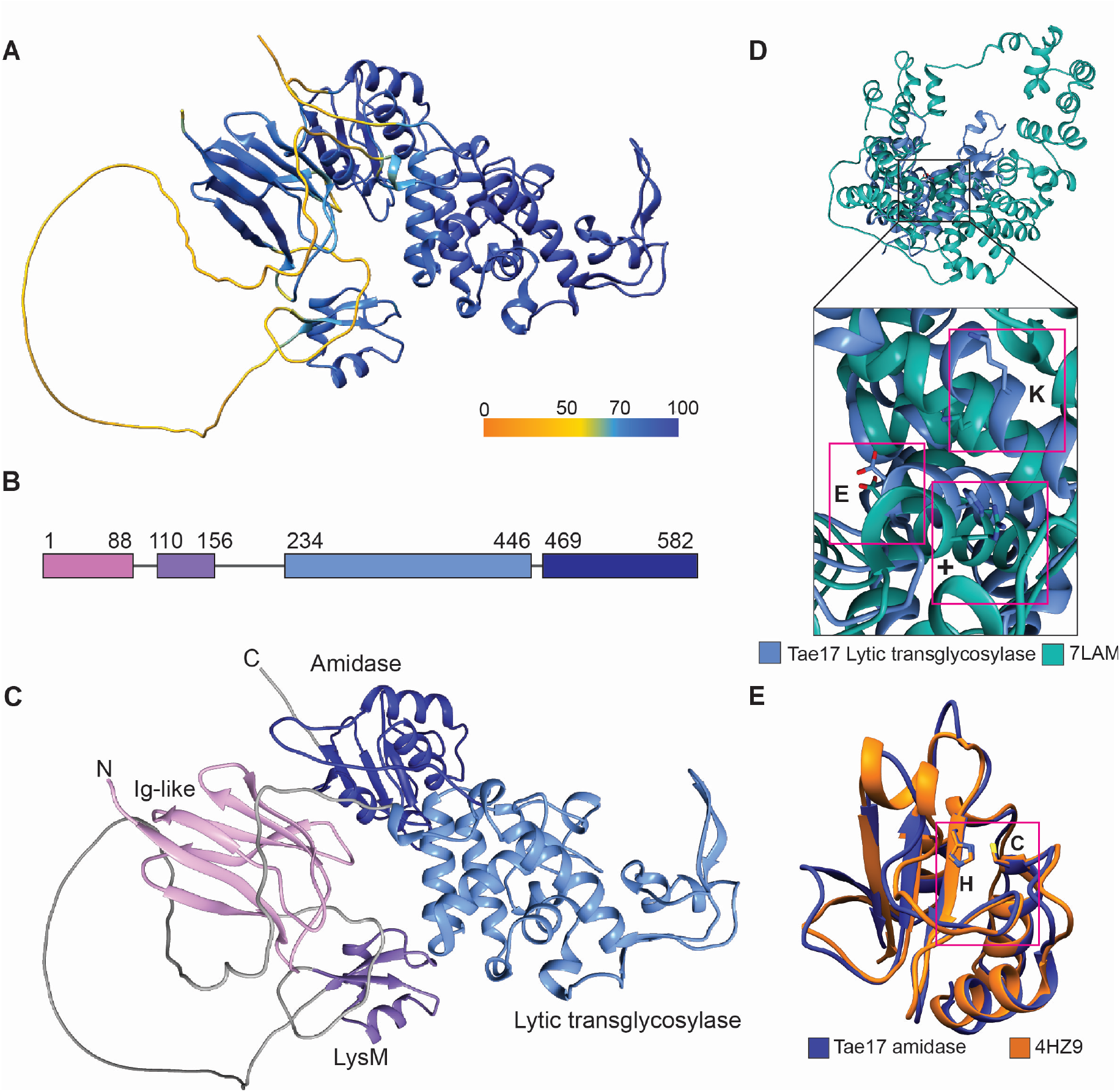
The Tae17 AlphaFold2 model has a four-domain structure with two likely active sites. (**A**) Tae17 AlphaFold2 model coloured according to confidence. (**B**) Tae17 sequence mapped by structural domains. Each domain is indicated in a different colour and residue numbers are shown. **(C)**Tae17 AlphaFold2 structure mapped by domains, colours are as used in panel B. (**D**) Alignment of the *C. jejuni* lytic transglycosylase 7LAM (green) and the proposed Tae17 lytic transglycosylase domain (light blue, residues 234-446). Active site resides are indicated by pink boxes: E278 (E390 in 7LAM), H276 (R388 in 7LAM; labelled +) and K436 (K505 for 7LAM). (**E**) Alignment of the proposed Tae17 amidase domain (dark blue, residues 469-582) to the full-length *Ralstonia pickettii* T6SS effector Tae3 (orange, PDB ID: 4HZ9). Active site residues are indicated by a pink box: C23 and H81 for Tae3 and the equivalent C488 and H550 for Tae17. For panel E, 4HZ9 chains B and C are removed as these are from the immunity protein Tai3.

To experimentally confirm that Tae17 had both lytic transglycosylase and amidase activity, we utilized the glucose-repressible and arabinose-inducible vector pBAD30 (30) to separately express wild-type and mutated Tae17 proteins in *E. coli* to assess their toxicity. As Tae17 is only toxic to *E. coli* when targeted to the periplasm (6), a sequence encoding a PelB leader was added to the 5’ end of each recombinant *tae17* gene. Expression plasmids encoding either wild-type Tae17 (Tae17^WT^), the lytic transglycosylase domain mutant Tae17^E278A^, the amidase domain mutant Tae17^C488A^, or a double mutant Tae17^E278A,C488A^, were separately expressed in *E. coli* from plasmids pAL1857, pAL1871, pAL1854 and pAL1870, respectively (Table 2).

**Table 1.**
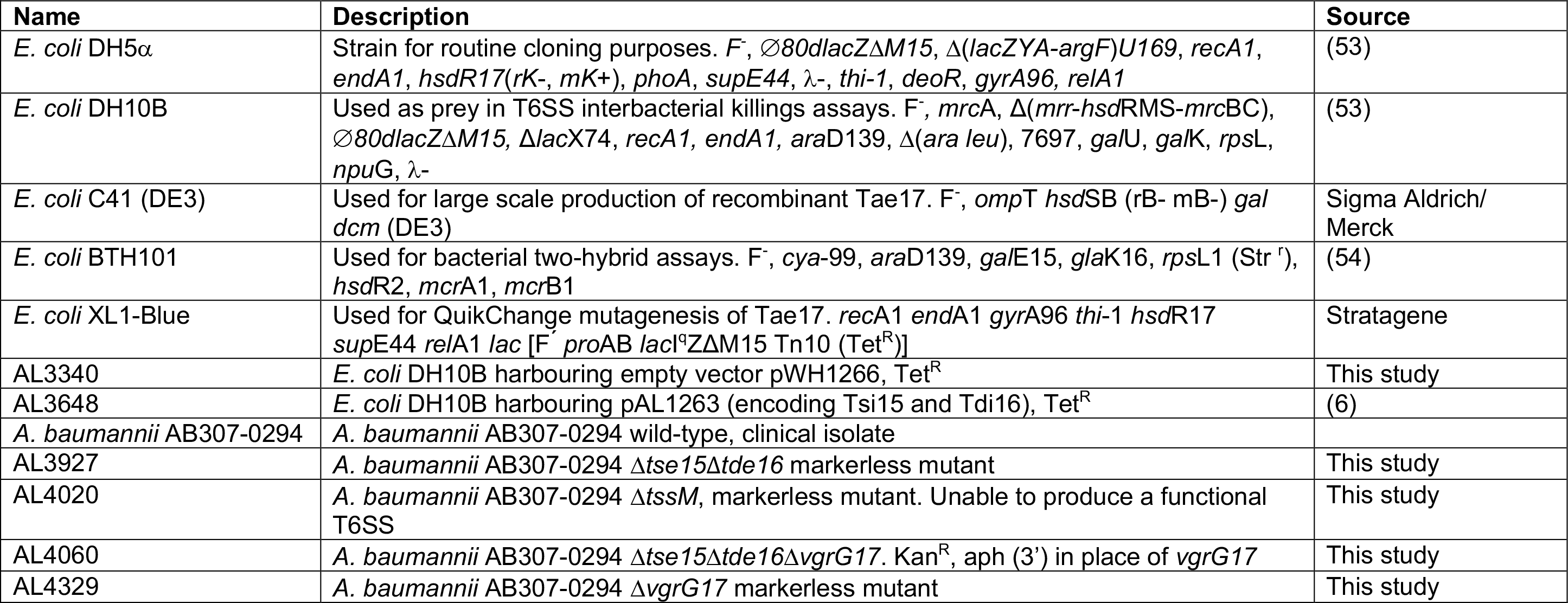
| Strains used in this study.

| Name | Description | Source |
| --- | --- | --- |
| *E. coli* DH5α | Strain for routine cloning purposes. *F*<sup>-</sup>, Δ80*dlacZ*ΔM15, $\Delta$(*lacZ*YA-*argF*)U169, *recA1*, *endA1*, *hsdR17*(*rK*<sup>-</sup>, *mK*<sup>+</sup>), *phoA*, *supE44*, λ<sup>-</sup>, *thi-1*, *deoR*, *gyrA96*, *relA1* | (53) |
| *E. coli* DH10B | Used as prey in T6SS interbacterial killings assays. *F*<sup>-</sup>, *mrcA*, $\Delta$(*mrr*-*hsdRMS*-*mrcBC*), Δ80*dlacZ*ΔM15, Δ*lacX74*, *recA1*, *endA1*, *araD139*, $\Delta$(*ara leu*), 7697, *galU*, *galK*, *rpsL*, *npuG*, λ<sup>-</sup> | (53) |
| *E. coli* C41 (DE3) | Used for large scale production of recombinant Tae17. *F*<sup>-</sup>, *ompT* *hsdSB* (*rB*<sup>-</sup> *mB*<sup>-</sup>) *gal dcm* (DE3) | Sigma Aldrich/ Merck |
| *E. coli* BTH101 | Used for bacterial two-hybrid assays. *F*<sup>-</sup>, *cya*-99, *araD139*, *galE15*, *glkA16*, *rpsL1* (Str<sup>r</sup>), *hsdR2*, *mcrA1*, *mcrB1* | (54) |
| *E. coli* XL1-Blue | Used for QuikChange mutagenesis of Tae17. *recA1* *endA1* *gyrA96* *thi-1* *hsdR17* *supE44* *relA1* *lac* [*F*<sup>-</sup> *proAB* *lacI*<sup>q</sup>ΔM15 Tn10 (Tet<sup>R</sup>)] | Stratagene |
| AL3340 | *E. coli* DH10B harbouring empty vector pWH1266, Tet<sup>R</sup> | This study |
| AL3648 | *E. coli* DH10B harbouring pAL1263 (encoding Tsi15 and Tdi16), Tet<sup>R</sup> | (6) |
| *A. baumannii* AB307-0294 | *A. baumannii* AB307-0294 wild-type, clinical isolate |  |
| AL3927 | *A. baumannii* AB307-0294 Δ*tse15*Δ*tde16* markerless mutant | This study |
| AL4020 | *A. baumannii* AB307-0294 Δ*tssM*, markerless mutant. Unable to produce a functional T6SS | This study |
| AL4060 | *A. baumannii* AB307-0294 Δ*tse15*Δ*tde16*Δ*vgrG17*. Kan<sup>R</sup>, *aph* (3') in place of *vgrG17* | This study |
| AL4329 | *A. baumannii* AB307-0294 Δ*vgrG17* markerless mutant | This study |

**Table 2.** | Plasmids used in this study.

| Name | Description | Source |
| --- | --- | --- |
| pAT03 | pMMB67EH encoding FLP recombinase that interacts with tandem FRT sites flanking kanamycin cassette (*aph* 3') to facilitate its removal. For generation of markerless *A. baumannii* mutants. Amp<sup>R</sup> | (47) |
| pBASE | *A. baumannii*/*E. coli* shuttle and recombinant protein expression vector (via the action of P<sub>tac</sub> promoter upstream of MCS). Amp<sup>R</sup> | (6) |
| pBAD30 | *E. coli* protein expression vector. Recombinant gene expression controlled P<sub>araBAD</sub> promoter (induced by arabinose and repressed by glucose). Amp<sup>R</sup> | (30) |
| pCR<sup>TM</sup>-Blunt II-TOPO<sup>TM</sup> | A vector used as template for the PCR amplification of the kanamycin resistance gene *aph* (3'). Gene used in SOE-PCR mediated mutagenesis of *A. baumannii* AB307-0294 | Invitrogen |
| pKT25 | B2H vector. Encoded the T25 fragment (1-224) of CyaA under control of a lac promoter. Kan<sup>R</sup> | (54) |
| pKT25-zip | B2H positive control plasmid. Kan<sup>R</sup> | (54) |
| pET28a | *E. coli* recombinant protein expression vector. Kan<sup>R</sup> | EMD Biosciences |
| pUT18C | B2H vector. Encodes the T18 fragment (amino acids 225 to 399) of CyaA under the control of the *lac* promoter. Amp<sup>R</sup> | (54) |
| pUT18C-zip | B2H positive control plasmid. Amp<sup>R</sup> | (54) |
| pWH1266 | *E. coli*/A. *baumannii* shuttle vector. Used in T6SS interbacterial killing assays to provide counter selection for *E. coli* DH10B. Tet<sup>R</sup>. | (55) |
| pAL1263 | pWH1266 encoding the T6SS immunity proteins Tsi15 and Tdi16, Tet<sup>R</sup>. Used in *E. coli* DH10B for counter selection (Tet<sup>R</sup>) and to provide immunity against the A. *baumannii* AB307_0294 T6SS effectors Tse15 and Tde16. | (6) |
| pAL1446 | pBASE encoding a truncated VgrG17 protein: amino acids 1-546. | (6) |
| pAL1447 | pBASE encoding the full-length VgrG17 protein (amino acids 1-1101) | (6) |
| pAL1565 | pBASE encoding truncated VgrG17: amino acids 1-1085 | This study |
| pAL1620 | pBASE encoding truncated VgrG17: amino acids 1-1074 | This study |
| pAL1621 | B2H plasmid pUT18C encoding VgrG17 amino acids 1001-1101 translationally fused to C terminal end of T18 | This study |
| pAL1623 | B2H plasmid pUT18C encoding VgrG17 amino acids 673-852 translationally fused to C terminal end of T18 | This study |
| pAL1624 | B2H plasmid pUT18C encoding VgrG17 amino acids 673-1101 translationally fused to C terminal end of T18 | This study |
| pAL1629 | B2H plasmid pKT25 encoding Tae17 amino acids 1-162 translationally fused to C terminal end of T25 | This study |
| pAL1630 | B2H plasmid pKT25 encoding Tae17 amino acids 155-485 translationally fused to C terminal end of T25 | This study |
| pAL1689 | B2H plasmid pKT25 encoding Tae17 amino acids 1-32 translationally fused to C terminal end of T25 | This study |
| pAL1699 | B2H plasmid pKT25 encoding Tae17 amino acids 33-162 translationally fused to C terminal end of T25 | This study |
| pAL1701 | B2H plasmid pKT25 encoding Tae17 amino acids 1-51 translationally fused to C terminal end of T25 | This study |
| pAL1702 | B2H plasmid pKT25 encoding Tae17 amino acids 14-63 translationally fused to C terminal end of T25 | This study |
| pAL1703 | B2H plasmid pUT18C encoding VgrG17 amino acids 1001-1050 translationally fused to C terminal end of T18 | This study |
| pAL1704 | B2H plasmid pUT18C encoding VgrG17 amino acids 1051-1085 translationally fused to C terminal end of T18 | This study |
| pAL1705 | B2H plasmid pUT18C encoding VgrG17 amino acids 1001-1085 translationally fused to C terminal end of T18. | This study |
| pAL1715 | B2H plasmid pKT25 encoding Tae17 amino acids 1-63 translationally fused to C terminal end of T25 | This study |
| pAL1716 | B2H plasmid pKT25 encoding Tae17 amino acids 1-116 translationally fused to C terminal end of T25 | This study |
| pAL1779 | pBASE containing *vgrG17* with a silent *SpeI* restriction site introduced for cloning purposes. Generated using SOE-PCR using products made from primer pairs BAP9260/BAP9262 and BAP9263/BAP9261 | This study |
| pAL1784 | pBASE encoding alanine substituted VgrG17<sup>G1069A,Y1070A</sup> | This study |
| pAL1785 | pBASE encoding alanine substituted VgrG17<sup>S1071A,A1072G</sup> | This study |
| pAL1786 | pBASE encoding alanine substituted VgrG17<sup>D1073A,K1074A</sup> | This study |
| pAL1787 | pBASE encoding alanine substituted VgrG17<sup>W1075A,H1076A</sup> | This study |
| pAL1789 | pET28a encoding a C-terminal His<sub>x6</sub>-tagged Tae17, codon-optimized for expression in *E. coli* | GenScript Biotech |
| pAL1793 | pBASE encoding alanine substituted VgrG17<sup>K1077A,E1078A</sup> | This study |
| pAL1794 | pBASE encoding alanine substituted VgrG17<sup>T1079A,E1080A</sup> | This study |
| pAL1795 | pBASE encoding alanine substituted VgrG17<sup>E1081A,F1082A</sup> | This study |
| pAL1796 | pBASE encoding alanine substituted VgrG17<sup>E1083A,D1084A</sup> | This study |
| pAL1797 | pBASE encoding alanine substituted VgrG17<sup>F1085A,D1086A</sup> | This study |
| pAL1809 | pBASE encoding alanine substituted VgrG17<sup>G1069A</sup> | This study |
| pAL1810 | pBASE encoding alanine substituted VgrG17<sup>Y1070A</sup> | This study |
| pAL1811 | pBASE encoding alanine substituted VgrG17<sup>W1075A</sup> | This study |
| pAL1812 | pBASE encoding alanine substituted VgrG17<sup>H1076A</sup> | This study |
| pAL1854 | pBAD30 encoding Tae17<sup>C488A</sup> with a PelB leader, codon-optimized for expression in *E. coli* | This study |
| pAL1857 | pBAD30 encoding Tae17<sup>WT</sup> with a PelB leader, codon-optimized for expression in *E. coli* | This study |
| pAL1870 | pBAD30 encoding Tae17<sup>E278A,C488A</sup> with a PelB leader, codon-optimized for expression in *E. coli* | This study |
| pAL1871 | pBAD30 encoding Tae17<sup>E278A</sup> with a PelB leader, codon-optimized for expression in *E. coli* | This study |
| pAL1885 | pET28a encoding a C-terminal His<sub>x6</sub>-tagged Tae17<sup>E278A,C488A</sup>, codon-optimized for expression in *E. coli* | GenScript Biotech |
| pAL1919 | pET28a encoding a C-terminal His<sub>x6</sub>-tagged Tae17<sup>E278A</sup>, codon-optimized for expression in *E. coli* | GenScript Biotech |
| pAL1920 | pET28a encoding a C-terminal His<sub>x6</sub>-tagged Tae17<sup>C488A</sup>, codon-optimized for expression in *E. coli* | GenScript Biotech |

To quantify the ability of each *E. coli* strain to survive heterologous expression of wild-type Tae17 or one of the Tae17 catalytic mutants, growth curves were performed, with or without induction of Tae17 expression. Both optical density (Fig. 2A) and cell viability (Fig. 2B) were assessed. All *E. coli* strains (expressing either recombinant Tae17^WT^, Tae17^E278A^, Tae17^C488A^ or Tae17^E278A,C488A^) grew normally when glucose was included in the medium to repress recombinant protein expression. Upon induction of protein expression, *E. coli* expressing Tae17^WT^ or the amidase mutant Tae17^C488A^ generated similar, low viability counts 5 h after induction (approximately 10^5^-fold lower than the corresponding uninduced strains). However, there was a small, but statistically significant, difference in the final optical density between the two cultures (Fig. 2A and B). *E. coli* cells expressing Tae17^E278A,C488A^ were as viable as the uninduced controls and grew at the same rate, showing that mutation of single active site residues in both predicted catalytic domains completely inactivated Tae17. An intermediate phenotype was observed when the expression of the lytic transglycosylase mutant Tae17^E278A^ was induced in *E. coli*. This strain showed reduced growth and viability when grown in the presence of arabinose but was not as attenuated for growth and viability as the strains expressing Tae17^WT^ or Tae17^C488A^ (Fig. 2A and B). Thus, under the conditions tested, both domains have lethal activity against *E. coli*; however, the predicted lytic transglycosylase domain appears to be more important for killing of *E. coli*.

**FIG 2.**
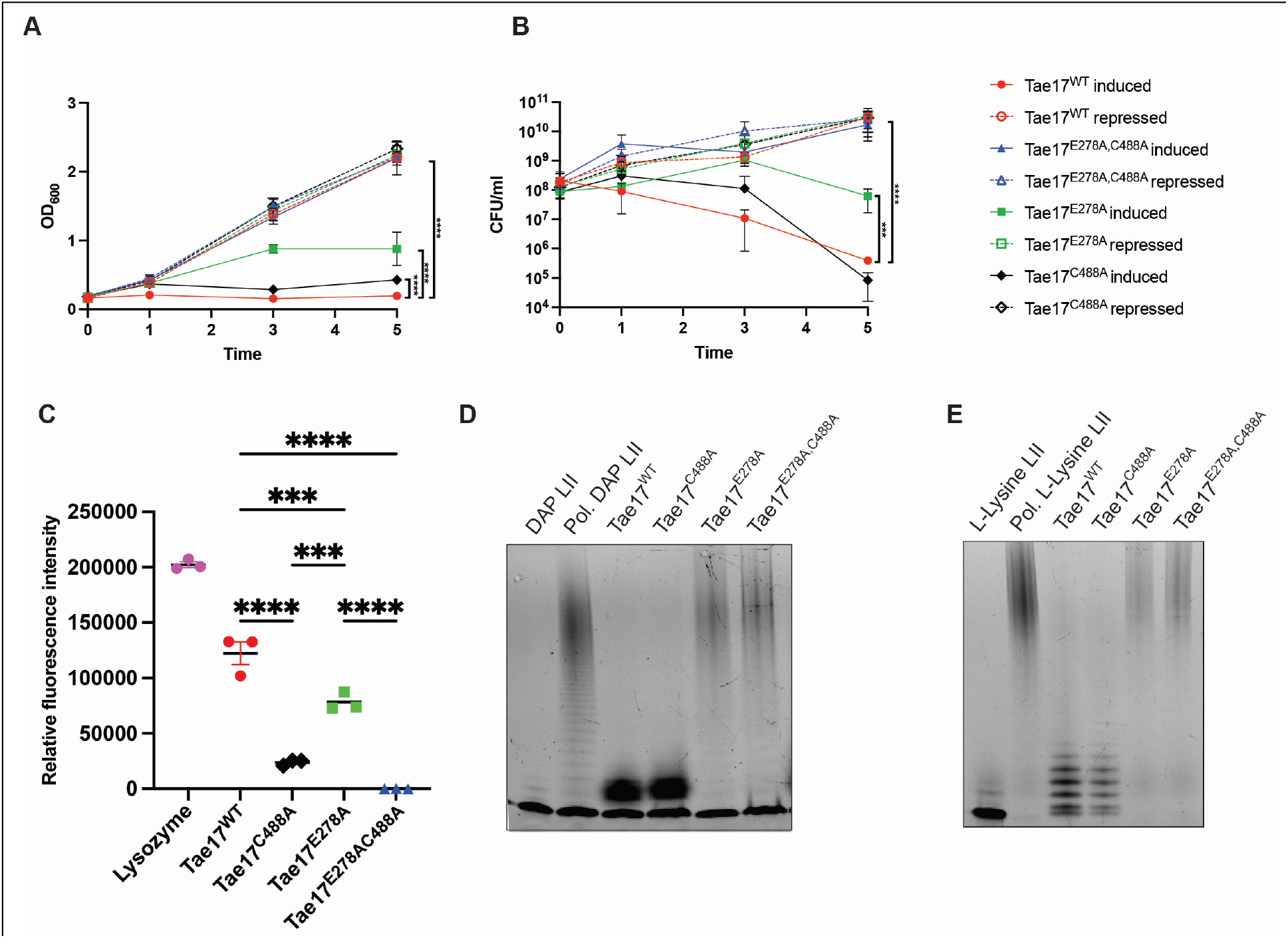
Activity of the *A. baumannii* AB307-0294 T6SS predicted bifunctional effector Tae17. (**A**) Growth curves were generated for recombinant *E. coli* strains harbouring a gene encoding Tae17^WT^, Tae17^E278A^, Tae17^C488A^, or Tae17^E278A,C488A^ (each with an added PelB leader sequence) on the vector pBAD30 (Table 2). Expression of the recombinant proteins was either induced with arabinose (solid markers and lines) or repressed with glucose (open markers and dotted lines) and optical density measured at 0 h, 1 h, 3 h and 5 h. (**B**) Viable counts were performed on the same cultures at 0 h, 1 h, 3 h and 5 h. On both graphs, each data point represents the mean of three biological replicates, and the vertical bars show ± SD. Statistical significance of the difference in means was analysed by ANOVA with Tukey’s multiple comparisons test. *** p<0.001, **** p<0.0001. (**C**) Degradation of FITC-labelled peptidoglycan following incubation for 2 h with purified lysozyme (control), Tae17^WT^, Tae17^C488A^, Tae17^E278A^, or Tae17^E278A,C488A^**(C)** An image of a Tris-Tricine gel following electrophoresis of the following samples: Gram-negative dansylated LII (DAP LII), polymerized DAP LII (Pol. DAP LII), and polymerized DAP LII following incubation for 2 h with purified Tae17^WT^, Tae17^C488A^, Tae17^E278A^ or Tae17^E278A,C488A^. (**E**) An image of a Tris-Tricine gel following electrophoresis of Gram-positive dansylated LII (L-Lysine LII), polymerized L- Lysine LII (Pol. L-Lysine LII), and polymerized L-Lysine LII following incubation for 2 h with purified Tae17^WT^, Tae17^C488A^, Tae17^E278A^ or Tae17^E278A,C488A^.

As most characterized T6SS peptidoglycan hydrolase effectors have only a single catalytic domain (4, 20, 28, 31) we were interested in further defining the function of these two domains in Tae17. Therefore, we purified recombinant Tae17^WT^, Tae17^E278A^, Tae17^C488A^ and Tae17^E278A,C488A^ from *E. coli* strains harbouring pAL1789, pAL1919, pAL1920 and pAL1885 respectively (Table 2), using nickel affinity-coupled size exclusion chromatography. For all constructs, the purified products resolved one predominant band at ∼67 kDa and a second band at ∼57 kDa on SDS-PAGE (Fig. S1). To determine the nature of the bands, we conducted peptide fingerprinting of Tae17^WT^. These data showed that the high MW species was indeed full-length Tae17 with 86% coverage, whereas the lower MW species was Tae17 with decreased peptide coverage prior to residue 91 (80% overall coverage) (data not shown). This suggests that for a proportion of the purified protein, the Ig-like domain (residues 1 – 88) had been cleaved. As this N-terminal degradation is unlikely to affect the catalytic activity of Tae17, we proceeded with functional analysis.

**Fig S1.**
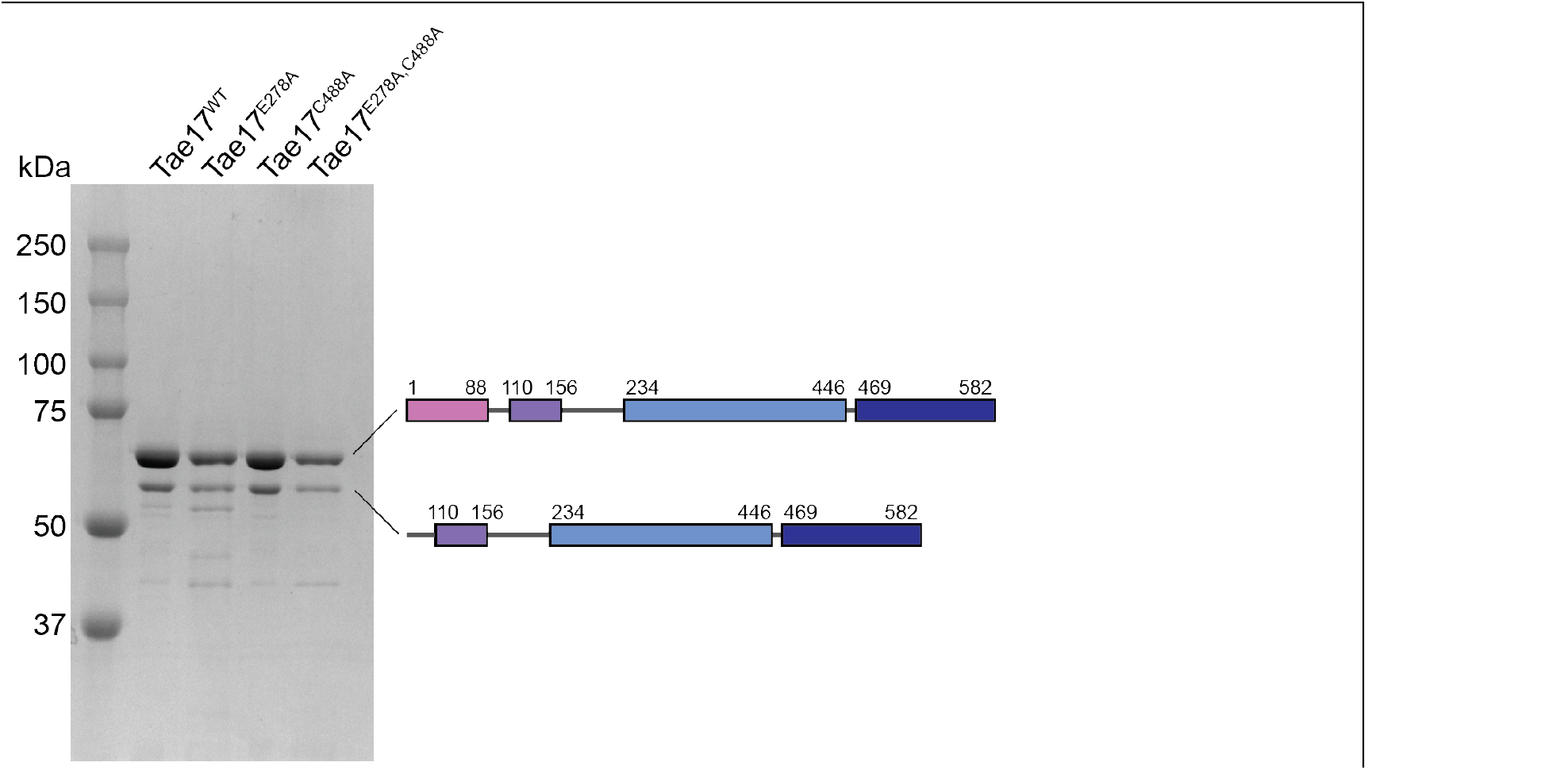
Purification of Tae17 and active site mutants. All proteins were purified by metal- affinity chromatography followed by size-exclusion chromatography. PAGE of Tae17, Tae17^WT^, Tae17^E278A^, Tae17^C488A^ and Tae17 ^E278AC488A^. The proposed domain structure of the two major species is shown at the right. Size markers are shown at the left.

To assess the ability of each of the purified proteins to degrade peptidoglycan, proteins were incubated with purified, intact FITC-labelled *E. coli* peptidoglycan and degradation activity measured. Incubation with Tae17^WT^ resulted in significant peptidoglycan degradation, while the activity of the dual catalytic mutant Tae17^E278A,C488A^ was undetectable. Both Tae17^C488A^ and Tae17^E278A^ showed significantly reduced but measurable peptidoglycan hydrolase activity. We then assessed their ability to degrade *in vitro* synthesized dansylated lipid II (LII) polymers that resembled either Gram-positive LII (L-Lys LII) or Gram-negative LII (DAP LII). Moreover, the Gram-positive representative L-Lys LII was amidated at position 2 of the peptide stem to iso-D-glutamine and had a lysine (Lys) at position 3 in the amino acid chain of peptidoglycan, which is representative of the chemical variant seen in *Staphylococcus aureus*, *Streptococcus pneumoniae* and other Gram-positive pathogens (32). The DAP LII, on the other hand, contained a diaminopimelic acid at position 3 (33). In both LII species, the presence of dansyl at DAP or Lys prevents transpeptidase crosslinking but allows for lipid II glycosyltransferase assembly of the glycan polymer. These polymers were separately incubated with the Tae17 recombinant proteins and the breakdown products generated were visualized on a Tris-Tricine gel (34), thus this assay is specific for lytic transglycosylase activity. Following the incubation of DAP LII with Tae17^WT^ or Tae17^C488A^, the DAP LII polymer was completely degraded (Fig. 2D), indicative of lytic transglycosylase activity. Incubation of DAP LII with Tae17^E278A^ or Tae17^E278A,C488A^ did not result in the breakdown of the DAP LII polymer, confirming that E278 is essential for the lytic transglycosylase activity. Both Tae17^WT^ and Tae17^C488A^ were also able to hydrolyse the glycosidic bonds of the L-Lys LII polymer, while Tae17^E278A^ and Tae17^E278A,C488A^ were not (Fig. 2E). This indicates that the lytic transglycosylase domain of Tae17 can hydrolyse the glycosidic linkage between MurNAc and GlcNAc in either the L-Lys LII or DAP LII polymers. Taken together, these data show that E278 is essential for the lytic transglycosylase activity against both Gram-positive and Gram-negative representative peptidoglycan, and while C488 is not involved in the breakdown of the glycosidic linkages, it is essential for the amidase activity.

### The N-terminus of Tae17 interacts with the C-terminus of the VgrG17 T6SS tip protein for effector delivery

We previously showed that the *A. baumannii* strain AB307_0294 lytic cargo effector Tae17 was crucial for interbacterial killing and dependent on VgrG17 for its delivery (6). However, the specifics of the interaction between the two proteins had not been defined. We therefore aimed to identify specific residues in VgrG17 that were required for interactions with Tae17. Previous studies have indicated that the C-terminal region of VgrG proteins confers specificity for their cognate effectors. Furthermore, Ig-like (or TTR-like) domains, ranging from of 8 to 62 amino acids in length, have been identified in this region and important for these interactions (15–17).

To determine if the C-terminal region of VgrG17 was involved in the interactions with, and T6SS-mediated delivery of, Tae17, we constructed expression plasmids (pAL1565 and pAL1620) encoding VgrG17 variants truncated at the C-terminal end by 16 amino acids (VgrG171-1085) and 27 amino acids (VgrG171-1074) (Fig. 3A). Each of the recombinant VgrG17 proteins was then tested for their ability to deliver Tae17 into *E. coli* using interbacterial killing assays. For these assays a triple mutant predator strain AB307-0294Δ*tse15*Δ*tde16*Δ*vgrG17* was constructed. This mutant lacks the T6SS effectors Tse15 and Tde16 that are also toxic to *E. coli* (6). Moreover, the mutant lacks a genomic copy of *vgrG17*. This predator strain can therefore only kill *E. coli* if a recombinant VgrG17 protein provided *in trans* interacts with and delivers the Tae17 effector. Interbacterial killing assays showed, as expected, that the AB307- 0294Δ*tse15*Δ*tde16*Δ*vgrG17* strain containing empty vector (EV) could not kill *E. coli* but providing the strain with wild-type VgrG17 *in trans* resulted in significant killing with *E. coli* viable counts below the level of detection (Fig. 3B). The AB307-0294Δ*tse15*Δ*tde16*Δ*vgrG17* strain expressing VgrG171-1085, lacking the C-terminal 16 amino acids, was still able to kill *E. coli* at similar levels as when the wild-type VgrG17 was expressed. However, the AB307- 0294Δ*tse15*Δ*tde16*Δ*vgrG17* strain expressing VgrG171-1074 could not kill *E. coli*. Therefore, these data indicate that one or more of the VgrG17 residues between amino acids 1075 and 1085 are crucial for the delivery of Tae17.

**FIG 3.**
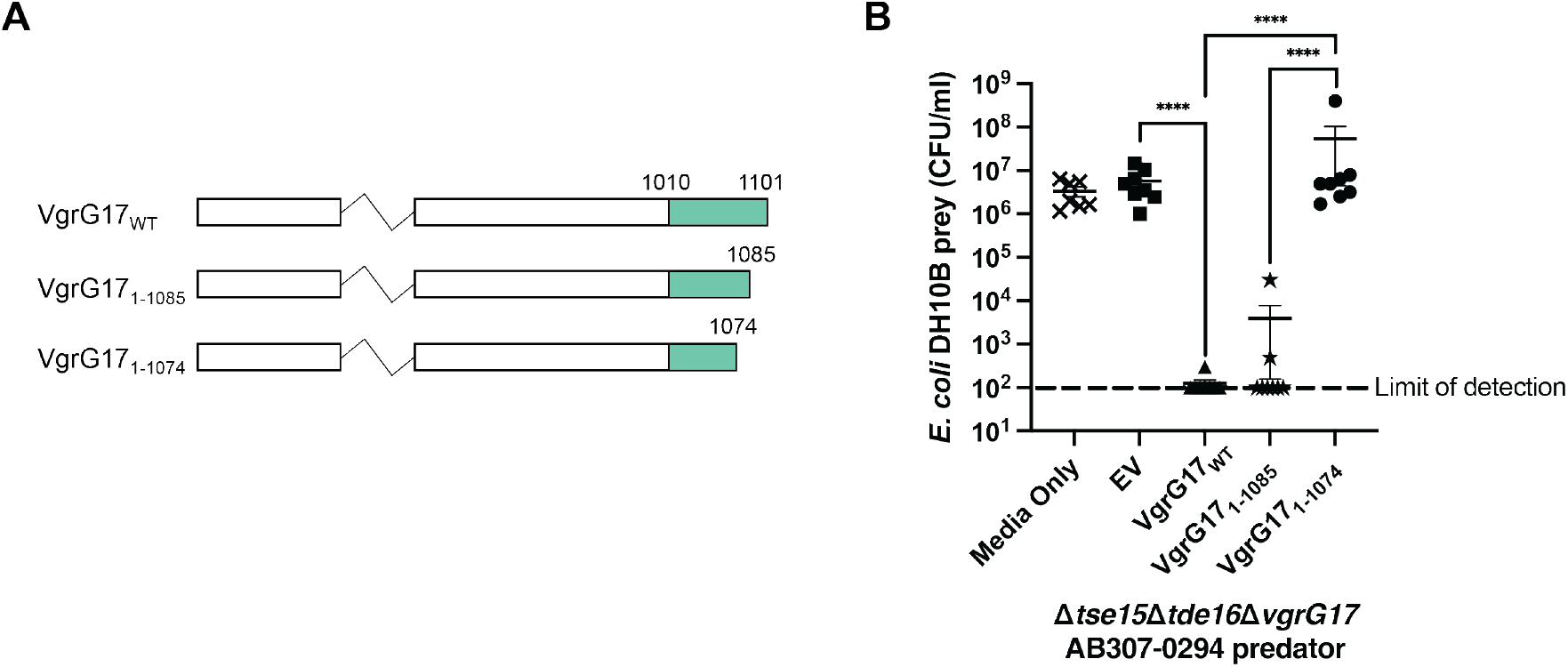
The effect of C-terminal truncations on the ability of VgrG17 to deliver Tae17 and kill *E. coli* prey. (**A**) A schematic representation of wild-type and C-terminal truncated VgrG17 proteins. The majority of *A. baumanni* VgrG sequence (white region) is conserved and predicted to participate in the formation of the stalk formed by the trimerization of VgrG proteins at the T6SS tip. This region is not shown in its entirety. The C-terminal end of the protein (green) is unique to VgrG17 and specific regions within it are predicted to interact with Tae17 for its delivery. (**B**) The ability of C-terminal truncated VgrG17 proteins VgrG171-1085 and VgrG171-1074 to deliver Tae17 via the T6SS was investigated using an interbacterial killing assay. The *A. baumannii* predator strain AB307- 0294τι*tse15*τι*tde16*τι*vgrG17* (strain AL4060) containing empty vector (EV), VgrG17, VgrG171-1085 or VgrG171-1074 was co-incubated for 3 h with *E. coli* prey strain DH10B harbouring an empty pWH1266 vector (AL3340). To check the growth of the prey in the absence of predator, the *E. coli* prey was also grown alone (Media Only). Each data point represents the mean of two technical replicates, the horizontal bars show the mean of the biological replicates, and the vertical bars show ± SEM. The dotted line signifies the limit of detection of the assay. Statistical significance of the difference in means was analysed by ANOVA with Tukey’s multiple comparisons test. n=7-8. **** p<0.0001.

To confirm there was a direct physical interaction between VgrG17 and Tae17, and identify the precise region involved in this interaction, we used bacterial adenylate cyclase two-hybrid (B2H) assays. A set of pUT18C-based plasmids, encoding the T18 bacterial adenylate cyclase fragment fused to fragments of VgrG17, and pKT25-based plasmids, encoding the T25 fragment fused to fragments of Tae17, were constructed and co-introduced into *E. coli* to assess the interaction between each recombinant protein (Table 2, Fig. S2A and S2B). Tae17 amino acids 1-162 (fused to the C terminal end of T25), were observed to interact with three T18-VgrG17 proteins, containing VgrG17 amino acids 1001-1101, 1001-1085 and 1051-1085 (Fig. S2C). These data show that the first 162 amino acids of Tae17 (containing the Ig-like domain and the LysM domain) interact with VgrG17 residues 1051-1085, which included the same region identified using interbacterial killing assays as important for interaction and delivery of Tae17 into *E. coli*.

**FIG S2.**
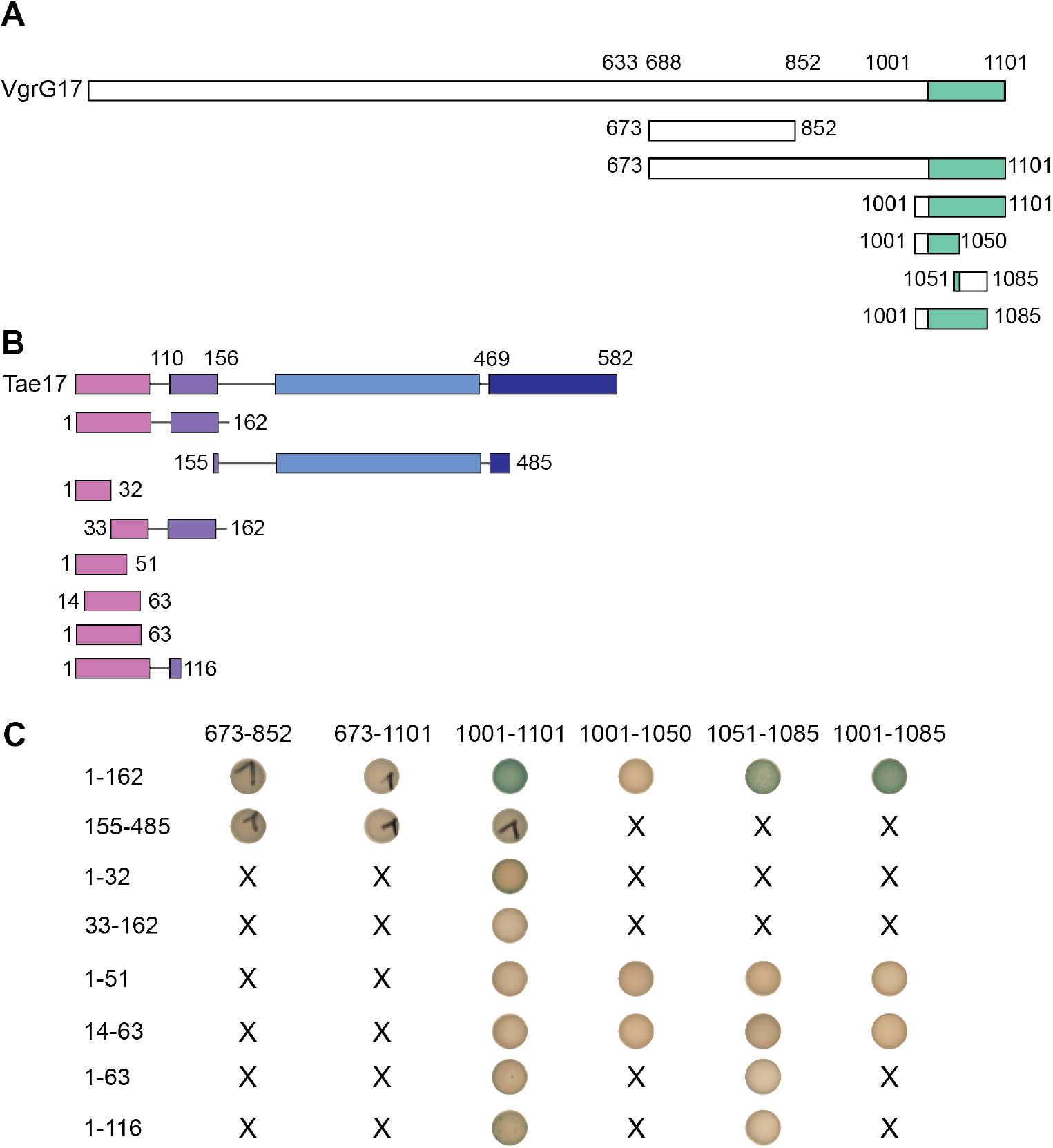
Bacterial adenylate cyclase two-hybrid analysis of the interacting regions of VgrG17 and Tae17. (**A**) Schematic representation of VgrG17 and the various regions used in the bacterial two-hybrid interaction studies. The white region indicates the conserved region of VgrG predicted to participate in the formation of the stalk formed by the trimerization of VgrG proteins at the T6SS tip. The C-terminal end of the protein (green) is unique to VgrG17 and specific regions within it are predicted to interact with Tae17 for its delivery. (**B**) A schematic representation of Tae17 and the regions that were assessed for their direct interaction with VgrG17 using bacterial adenylate two hybrid experiments. Domains are coloured according to structure in Figure 1. (**C**) Figure showing T18-VgrG17 and T25-Tae17 proteins tested for their interaction. Numbers on the left indicate region of Tae17 fused to T25 adenylate cyclase fragment. Numbers at top of figure indicate region of VgrG17 fused to the T18 fragment of adenylate cyclase. The presence of interaction between pairs of proteins in the assay is indicated by the formation of blue colonies on selective media (actual colony images are shown). Text crosses indicate VgrG17 and Tae17 region pairs that were not tested.

### VgrG17^G1069^ and VgrG17^W1075^ are important for the delivery of Tae17

The interbacterial killing assays (above) indicated that one or more amino acids between amino acids 1075 and 1085 in VgrG17 was required to facilitate Tae17-mediated killing of *E. coli*. The importance of this region was supported by the B2H data, which indicated that the region in VgrG17 required for interaction with Tae17 was within amino acids 1051-1085. To determine which of the specific residues in this region are most important for this interaction and subsequent delivery of the Tae17 effector, we used alanine scanning mutagenesis. We first made a series of VgrG17 expression plasmids using the *A. baumannii* pBASE expression vector, each encoding full-length VgrG17 but with specific dual amino acid substitutions between residues 1069-1086 (Fig. 4A). In total, nine proteins were separately expressed *in trans* in the AB307-0294*Δtse15Δtde16ΔvgrG17* predator strain and each strain assessed for the ability to kill *E. coli* DH10B prey cells in a Tae17-dependent manner. Killing activity was initially assessed qualitatively by plating all co-incubation growth onto LB agar with tetracycline that allowed only *E. coli* survivors to grow. After 3 h co-incubation, only two of the nine *A. baumannii* predator strains failed to kill *E. coli* (data not shown), indicating that the expressed recombinant proteins did not interact with, and deliver, Tae17. These strains expressed, *in trans*, VgrG17^G1069A,Y1070A^ or VgrG17^W1075A,H1076A^, suggesting that one or both of residues G1069 and Y1070, and W1075 and H1076 were essential for delivery of Tae17.

**FIG 4.**
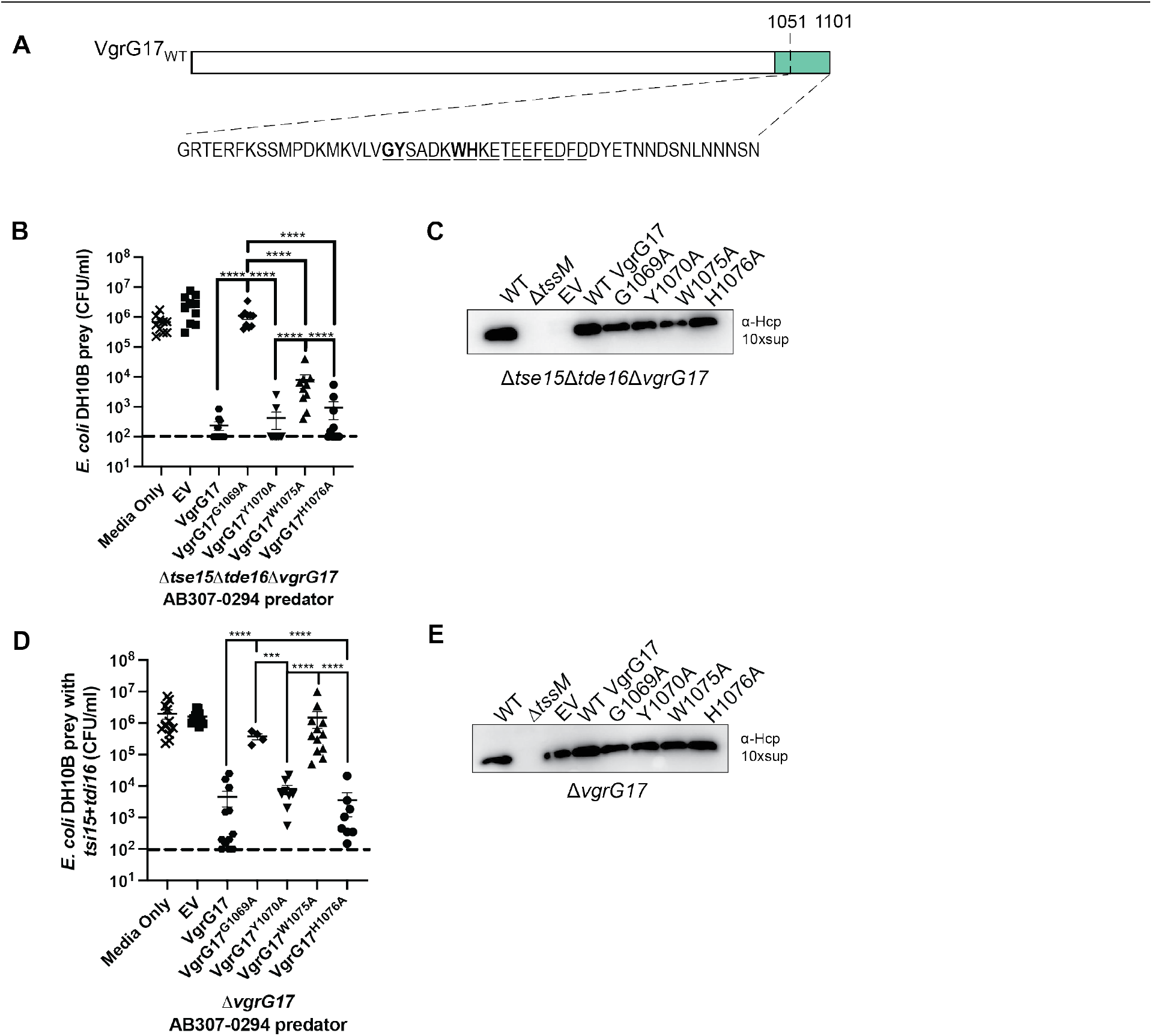
VgrG17 amino acids G1069 and W1075 are important for delivery of the effector Tae17. (**A**) Schematic representation of VgrG17 with amino acid sequence in region of interest (1051–1101) shown. Amino acids targeted for dual alanine substitutions are shown in underlined pairs while amino acids targeted by single alanine substitutions (G1069, Y1070, W1075 and H1076) are shown in bold. The white region of VgrG indicates the portion predicted to participate in the formation of the stalk formed by the trimerization of VgrG proteins at the T6SS tip, while the region in green is predicted to be the general region available for interaction with Tae17. (**B**) Interbacterial killing assay using *E. coli* DH10B harbouring empty pWH1266 vector as prey, co-cultured for 3 h with the *A. baumannii* predator strain AB307- 0294τι*tse15*τι*tde16*τι*vgrG17* (AL4060) as predator, provided with empty pBASE (EV) or a pBASE- derivative encoding wild-type VgrG17, or VgrG17 with a single amino substitution as shown. As a prey growth control, *E. coli* DH10B harbouring empty pWH1266 was cultured alone (Media Only) (**C**) Western blot assessing Hcp secretion in the *A. baumannii* predator strain AB307-0294τι*tse15*τι*tde16*τι*vgrG17* provided with plasmids encoding wild-type VgrG17 or one of the VgrG17 alanine mutants. The secretion of the T6SS needle protein, Hcp, is used as an indicator of a functional T6SS in *A. baumannii*. (**D**) Interbacterial killing assay using *E. coli* DH10B harbouring a plasmid encoding immunity proteins Tdi16 and Tsi15 as prey (to neutralize T6SS effectors Tse15 and Tde16). The *A. baumannii* predator strain in this assay was AB307-0294τι*vgrG17* (AL4329) containing EV, or a pBASE-derivative encoding wild-type VgrG17, or VgrG17 with a single amino substitution as shown. (**E**) Western blot assessing Hcp secretion by the AB307-0294τι*vgrG17* predator strain provided with plasmids encoding wild type VgrG17 or one of the VgrG17 alanine mutants. For panels A and C, each data point represents the mean of two technical replicates, the horizontal bars show the mean of the biological replicates, and the vertical bars show ± SEM. The dotted line signifies the limit of detection of the assay. Statistical significance of the difference in means was analysed by ANOVA with Tukey’s multiple comparisons test. n=4-12. *** p=0.0001, **** p<0.0001. For panels C and E, an AB307-0294 τι*tssM* strain was used as a negative control as it has a non-functional T6SS. All supernatants were concentrated 10x (10xsup).

To determine which of the four VgrG17 amino acids identified in the dual alanine mutagenesis qualitative assays were important for interaction with Tae17, we generated expression plasmids (again using the *A. baumannii* expression plasmid pBASE) encoding the following single mutants: VgrG17^G1069A^, VgrG17^Y1070A^, VgrG17^W1075A^ and VgrG17^H1076A^ (Fig. 4A). We then used quantitative interbacterial killing assays to assess which of the alanine substituted VgrG17 proteins was still able to interact with and deliver Tae17 to kill *E. coli*. The separate expression of VgrG17^Y1070A^ and VgrG17^H1076A^ in the AB307-0294Δ*tse15*Δ*tde16*Δ*vgrG17* predator strain fully restored the ability of *A. baumannii* to kill *E. coli* via Tae17 activity, indicating that these residues alone are not crucial for interaction with Tae17 (Fig. 4B). However, the predator cells expressing VgrG17^G1069A^ were unable to kill *E. coli*. Thus, amino acid G1069 in VgrG17 appears essential for the interaction with Tae17. The predator strain expressing VgrG17^W1075A^ also showed significantly reduced killing, suggesting that amino acid W1075 in VgrG17 is also important for the interaction with Tae17.

To assess whether the single alanine substitutions in VgrG17 were specifically affecting Tae17 binding and delivery, and not abrogating the formation or activity of the T6SS apparatus itself, we used the secretion of the needle protein Hcp as a marker for T6SS assembly and activity. We assessed the ability of the AB307-0294Δ*tse15*Δ*tde16*Δ*vgrG17* strains, each provided with a pBASE plasmid encoding a different alanine-substituted VgrG17 protein, to secrete Hcp into the supernatant (Fig. 4C). Interestingly, the AB307-0294Δ*tse15*Δ*tde16*Δ*vgrG17* strain harbouring empty pBASE vector had an inactive T6SS with no observable Hcp secretion. This indicates that the AB307-0294 T6SS lacking VgrG17 does not function in the absence of other functional VgrG/effector pairs; similar results have been observed previously in other species (35). However, providing the AB307-0294τι*tse15*τι*tde16*τι*vgrG17* strain with wild-type VgrG17 or any of the single VgrG17 alanine mutants *in trans* restored Hcp secretion (Fig. 4C). This included the AB307-0294τι*tse15*τι*tde16*τι*vgrG17* strain expressing VgrG17^G1069A^, which our interbacterial killing assay showed cannot deliver Tae17. Together, the data suggest that in the absence of Tse15 and Tde16 effector proteins, a full-length VgrG17 protein is essential for T6SS function, even if the VgrG17 protein supplied to the strain, e.g. VgrG17^G1069A^, cannot bind and deliver the Tae17 effector. These data also show that complementation with any of the mutant VgrG17 proteins can form a fully active system, which indicates that each of these mutated VgrG17 proteins can form a functional VgrG trimer and T6SS structure.

To confirm the interbacterial killing results obtained using the AB307- 0294τι*tse15*τι*tde16*τι*vgrG17* predator strain, which showed no T6SS activity in the absence of a functional VgrG17, we performed additional interbacterial killing assays using a predator strain that produced all three T6SS effectors, Tse15, Tde16 and Tae17. The AB307- 0294τι*vgrG17* mutant (AL4329) can deliver the effectors Tse15 and Tde16 (via VgrG15 and VgrG16, respectively) but can only deliver Tae17 when a functional VgrG17 protein is expressed *in trans* from the complementing plasmid. To allow the use of this predator strain for specifically assessing Tae17 delivery, the *E. coli* prey strain was provided with the appropriate cognate immunity proteins to protect against the T6SS effectors Tse15 and Tde16 (Tsi15 and Tdi16, respectively). In this case, any T6SS-mediated killing of *E. coli* by AB307- 0294τι*vgrG17* can be attributed directly to successful VgrG17-mediated delivery of Tae17, as has been previously shown (6). As expected, in these interbacterial killing assays the *in trans* expression of wild-type VgrG17, VgrG17^Y1070A^ or VgrG17^H1076A^ fully restored the ability of the AB307-0294τι*vgrG17* predator strain to kill *E. coli* via Tae17 delivery (Fig. 4D). However, when plasmids expressing either VgrG17^G1069A^ or VgrG17^W1075A^ were provided to AB307- 0294τι*vgrG17*, the strains were unable to kill *E. coli* (Fig. 4D). These data confirm that the VgrG17 amino acids G1069 and W1075 are crucial for interaction with, and delivery of, Tae17. To check the activity of the T6SS in these strains, we measured Hcp secretion as described earlier. All AB307-0294 τι*vgrG17* strains harbouring empty vector or a VgrG17 expression plasmid produced similar levels of secreted Hcp, thereby confirming the T6SS was active in this mutant with or without provision of a functional VgrG17 (Fig. 4E).

### Tae17 and VgrG17 are predicted to interact via edge-to-edge contacts across two domains

The VgrG17 experimental data obtained using truncated and alanine substituted proteins and B2H data, indicated that the VgrG17 region from 1051-1085 is necessary for interaction with Tae17, and that the VgrG17 amino acids G1069 and W1075 are required for this interaction. Furthermore, for Tae17, the B2H data indicated that the first 162 amino acids are important for the interaction with this region of VgrG17. To understand the likely molecular basis of these interactions, we used AlphaFold2 multimer to model the interaction of Tae17 with this region of VgrG17.

The T6SS needle tip is composed of a trimeric VgrG protein complex, where the N-terminal and central sections of each VgrG protein interact to form a stalk that is held together by strong intermolecular interactions (13). However, existing sequence and structural data indicate that the C-terminal regions of the VgrG proteins do not interact with each other and are free to interact with the effectors (13). Therefore, to identify potential contacts and align this to our experimental data, we modeled the C-terminal amino acids 1010-1101 of VgrG17, which fully encompassed the region shown to be important for VgrG17:Tae17 interaction, with full-length Tae17. Initial inspection identified two regions of interacting β-strands. One of these regions includes residues 1013-1070 of VgrG17, which fold into two small β-sheets that together form a sandwich-like structure that interacts with a β-strand formed by residues 90-93 of Tae17 (Fig. 5). We note the structural and functional similarity of this sub-structure to that identified by Flaugnatti *et al.* (36), and hence, we term this the transthyretin-like (TTR) domain. One sheet of this TTR comprises two β-strands from VgrG17, while the other sheet comprises three β-strands from VgrG17 and a single β-strand from Tae17. The second of the interacting β- strand regions included residues 1081-1083 of VgrG17 and a six-stranded Ig-like domain formed by two β-sheets of Tae17 from residues 4-93 (Fig. 5). Overall, the C-terminal region of VgrG17 (from residues ∼1071-1084) appears to thread through the multi-domain structure of Tae17 toward the amidase domain (Fig. S3). The C-terminal ∼17 amino acids of VgrG17 (residues 1084-1101) are modeled as an α-helix that does not interact with Tae17, which supports our VgrG17 C-terminal truncation data that this region is not involved in the interaction (Fig. 3).

**FIG 5.**
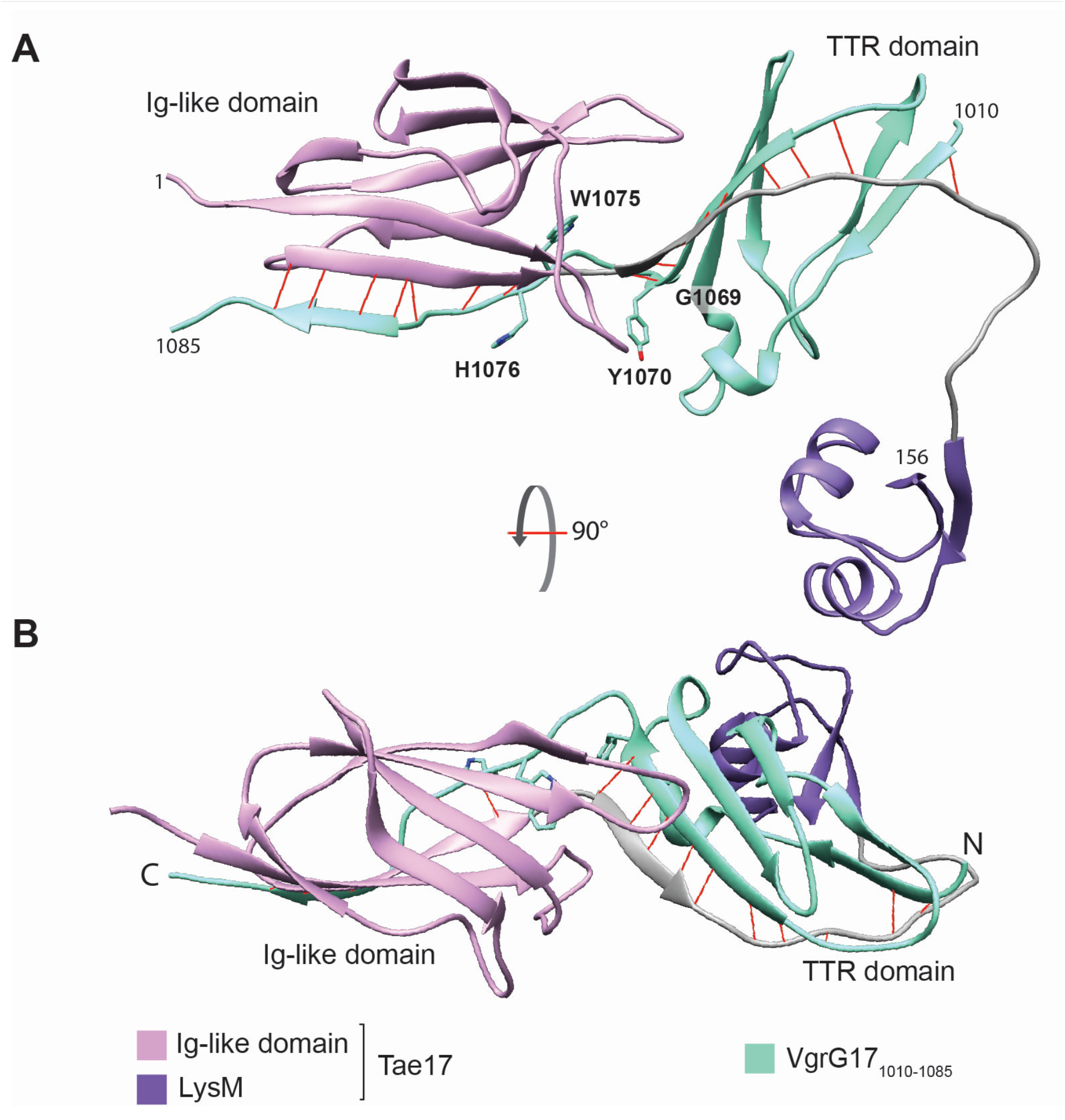
(**A**) AlphaFold2 model of VgrG171010-1085 interacting with Tae171-156. While the AlphaFold2 interaction was modelled with the full-length Tae17, only the interacting region is shown here for clarity. VgrG17 is shown in green. For Tae17, the Ig-like domain is shown in pink, the LysM domain is shown in purple and the linker region is coloured grey. Edge-to- edge contact backbone hydrogen bonds are shown in red. Residues of interest mutated during the alanine scanning mutagenesis are shown. The amidase and lytic transglycosylase domains are removed for clarity but are included in FIG S3. (**B**) The same AlphaFold2 model of VgrG171010-1085 interacting with Tae171-156 but rotated forward 90 degrees. The VgrG17 N- and C-termini and TTR domain are indicated.

To look more specifically at how VgrG17 and Tae17 interact in the model, the coordinates were visually inspected and input into PISA (37). No disulfide or covalent bonds were present, consistent with what is known about cargo effectors. However, approximately 17 backbone hydrogen bonds between the edge-to-edge contacts were predicted (Fig. 5 in red) as well as up to 12 salt bridges (Table S1, Fig. S3, Fig. S4, Movie S1). The large number of salt bridges predicted across the interface is due to the highly negatively charged VgrG17 C-terminus, and highly positively charged Tae17 N-terminus. This suggests that electrostatics are likely important for the interaction between VgrG17 and Tae17. Notably, a number of these charged residues were mutated during the dual alanine scanning (D1073, E1078, E1080, E1081, E1083, D1084, D1086) yet these mutations showed no change in the delivery of Tae17. Hence, Tae17 can deliver VgrG17 even when some of the interacting residues have been mutated. As charged interactions are mediated by overall electrostatic potential, individual or dual mutation of these charged residues likely does not reduce the overall electrostatic potential enough to prevent the interaction.

**Table S1.**
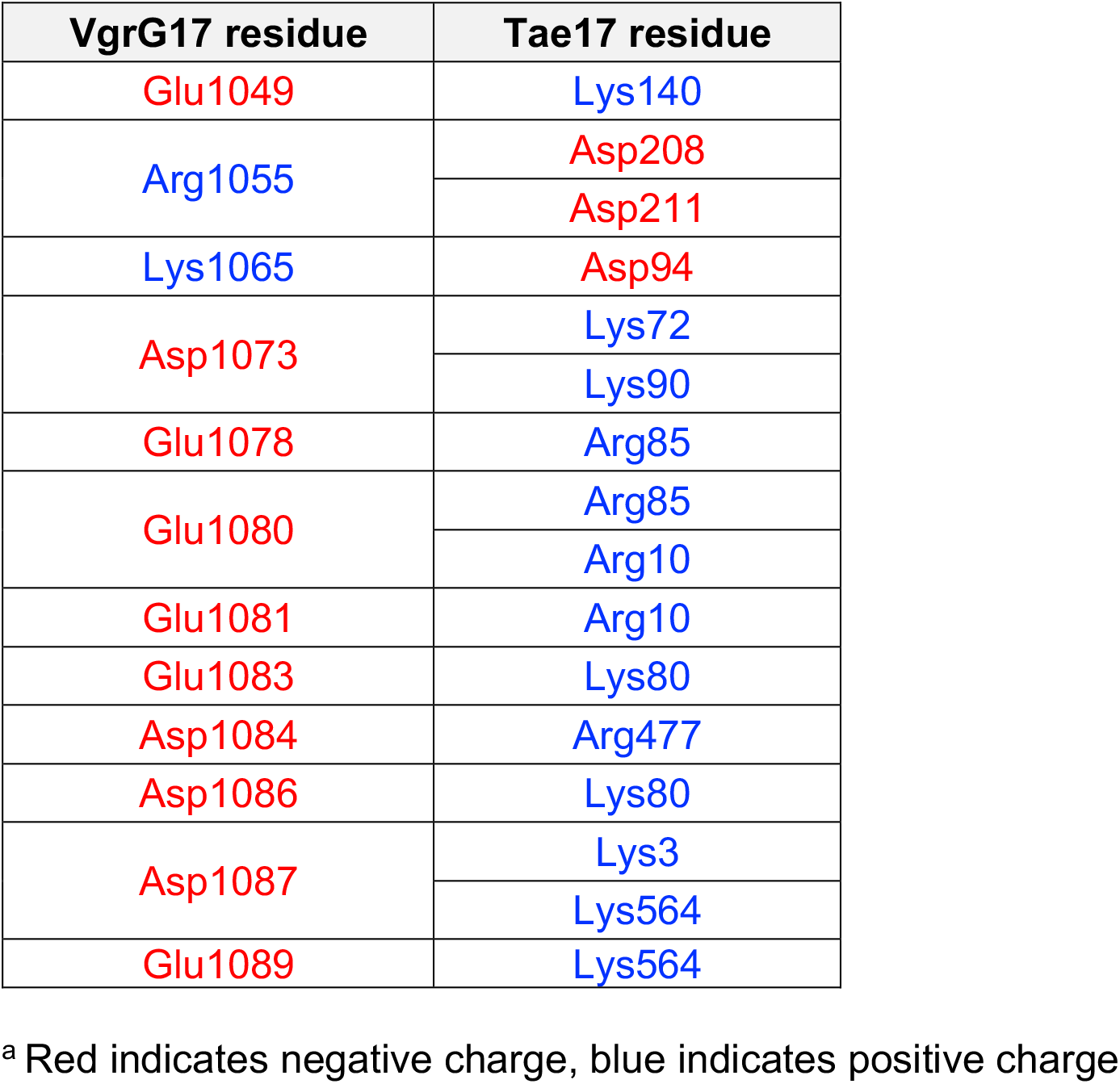
VgrG17:Tae17 AlphaFold2 model salt bridges.

| VgrG17 residue | Tae17 residue |
| --- | --- |
| Glu1049 | Lys140 |
| Arg1055 | Asp208 |
|  | Asp211 |
| Lys1065 | Asp94 |
| Asp1073 | Lys72 |
|  | Lys90 |
| Glu1078 | Arg85 |
| Glu1080 | Arg85 |
|  | Arg10 |
| Glu1081 | Arg10 |
| Glu1083 | Lys80 |
| Asp1084 | Arg477 |
| Asp1086 | Lys80 |
| Asp1087 | Lys3 |
|  | Lys564 |
| Glu1089 | Lys564 |

<sup>a</sup> Red indicates negative charge, blue indicates positive charge

**FIG S3.**
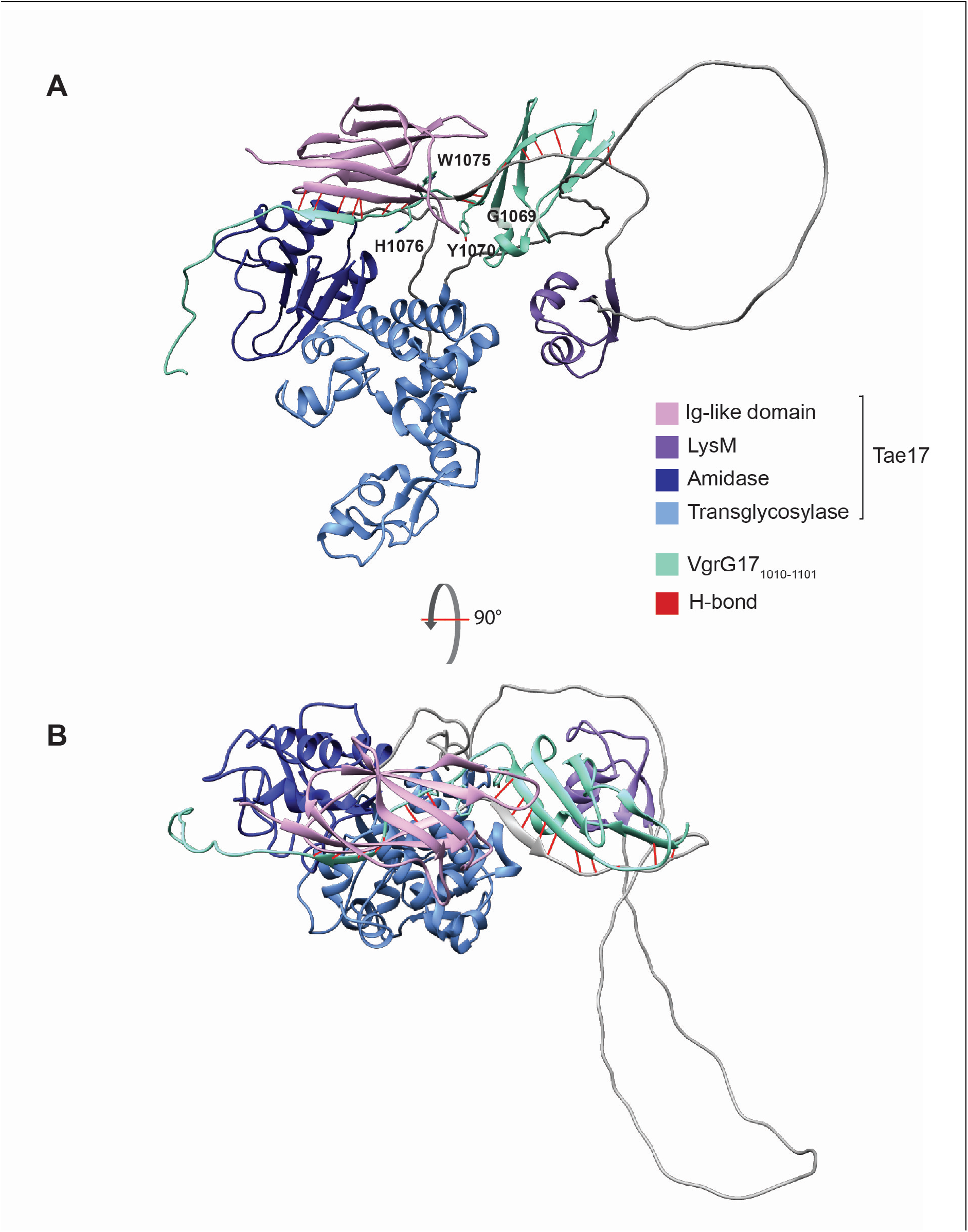
(A) AlphaFold2 model of VgrG171010-1101 interacting with full length Tae17. VgrG17 is shown in green. The Tae17 Ig-like domain is shown in pink, the LysM domain is shown in purple, lytic transglycosylase domain in light blue and amidase domain in dark blue. The linker regions are coloured grey. Edge-to-edge contact hydrogen bonds are shown in red. Residues of interest mutated during the alanine scanning mutagenesis are shown. (B) The same VgrG171010-1101 interaction with full length Tae17 but rotated forward 90 degrees.

**FIG S4.**
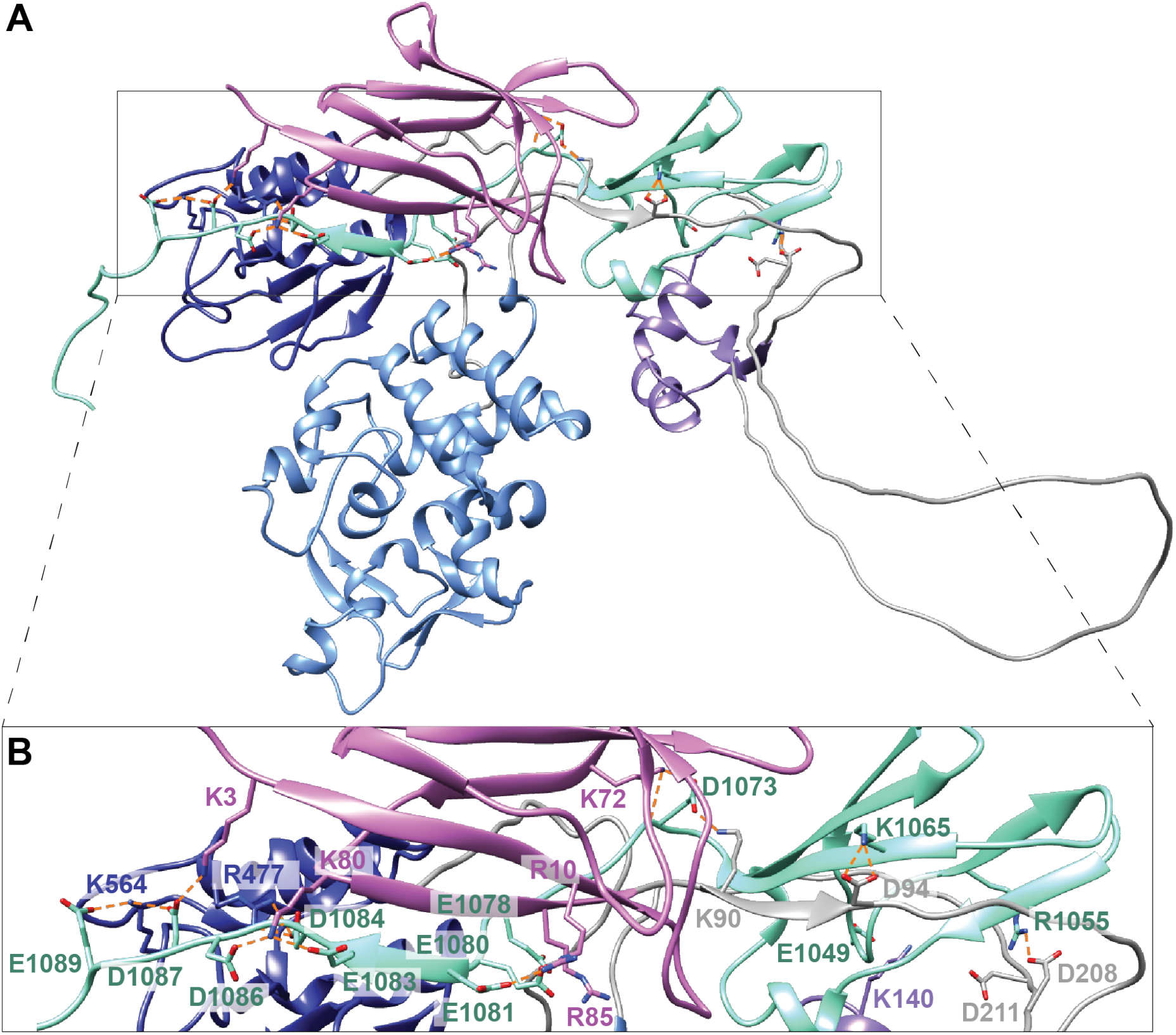
AlphaFold2 model of VgrG171010-1101 interacting with Tae17 with predicted salt bridges shown. For both **A**) and **B**) VgrG17 is shown in green, Tae17 Ig-like domain is shown in pink, the LysM domain is shown in purple, lytic transglycosylase domain in light blue and amidase domain in dark blue. The linker regions are coloured grey. All residues predicted to form salt bridges by PISA are shown as sticks and further coloured by heteroatom. All salt bridges that can form between these residues, as determined by Chimera, are shown as dashed orange lines. All other PISA predicted salt bridges are possible following minor conformational changes, as such we have shown all residues even if bonds were not predicted using Chimera. **A**) shows the full length Tae17 and VgrG171010-1101 with salt bridges. **B**) depicts a zoomed version of panel A) where each of the residues is also labelled using single letter code. The colour of the label identifies the chain or domain that the residue is found within. For further visualization of these interactions, please see the supplementary movie.

## DISCUSSION

Numerous Gram-negative bacteria employ a T6SS to deliver antibacterial effector proteins into neighbouring bacterial cells as a means of gaining dominance in polymicrobial environments. The majority of bacteria that possess a T6SS, including *A. baumannii*, are human, animal or plant pathogens (38). Consequently, manipulating the T6SS could be an important novel strategy for control of pathogens such as *A. baumannii, Pseudomonas aeruginosa, Vibrio cholerae* and *Serratia marcescens* (39–41). Recent bioinformatic analysis of T6SS effectors in *A. baumannii* strains identified >30 phylogenetically distinct families (23), with at least eight of these predicted to be involved in the degradation of peptidoglycan. The amino acid sequences of these effectors are very diverse, suggesting that these different effector families will have unique mechanisms of action. This study aimed to explore the structure, function and delivery determinants of a group 17 peptidoglycan hydrolase, Tae17, from the *A. baumannii* clinical strain AB307-0294.

AlphaFold2 modelling of Tae17 revealed a modular protein with four domains separated by linkers. This analysis suggested that Tae17 contains two catalytic domains: one with lytic transglycosylase activity and one with amidase activity. Mutation of the proposed active site residues in these domains, alone and in combination, confirmed that Tae17 is indeed a bifunctional peptidoglycan hydrolase. Tse4 from *A. baumannii* ATCC 17978 has also recently been identified as a bifunctional peptidoglycan hydrolase (21) but belongs to a different effector family and shares no readily identifiable amino acid sequence identity with Tae17 (23). Previous research on other amidase proteins, including Tae3 (28), Tse1/Tae1 (42) and Tae4 (43), has demonstrated that mutation of a single catalytic residue can completely abolish effector activity. However, mutation of the Tae17 predicted amidase domain alone did not impede the ability of the effector to kill *E. coli.* This suggests that under these conditions, the lytic transglycosylase domain is the dominant domain of Tae17, and that *E. coli* may not be the primary target of the Tae17 amidase domain.

Gram-positive and Gram-negative bacteria produce peptidoglycan that is chemically different. Both contain a similar backbone of repeating MurNAc and GlcNAc sugar units, but different stem peptide chemical components are observed. We tested Tae17 activity against *in vitro* synthesized Gram-positive LII-type or Gram-negative LII-type peptidoglycan backbone. The Tae17 proteins with an intact lytic transglycosylase domain were able to degrade both types of *in vitro* synthesized peptidoglycan. Therefore, this domain can recognize and cleave the glycosidic bonds in the peptidoglycan sugar chain that is representative of both Gram-positive and Gram-negative peptidoglycan (33).

For the Tae17 effector to be delivered, it must interact with the structural components of the T6SS. Our previous study showed that Tae17 specifically interacts with VgrG17 for delivery (6). In this current study, we have shown that the N-terminal 162 amino acids of Tae17 interact directly with the C-terminal region of VgrG17. Deletions from either end of this short Tae17 fragment resulted in the loss of interaction with VgrG17, strongly suggesting that interaction required both the Ig-like domain and LysM domain of Tae17. We also identified the region of VgrG17 between residues 1051-1085 as important for interaction with Tae17, and specifically residues G1069 and W1075, as measured by interbacterial killing assays. These data support and significantly extend a number of previous studies that show the effector N-terminal region interacts with the C-terminal region of VgrG for delivery by the T6SS (15) (16) (17). Our data present, for the first time, residues essential for the interaction required for T6SS cargo effector delivery.

Utilising the experimental interaction data generated in this study, we produced an AlphaFold2 model of the Tae17:VgrG17 interaction. These data predicted two edge-to-edge contact interactions between two different domains, reminiscent of a related experimentally determined structure (36). One interaction involved the Ig-like domain of Tae17, and a single short β-strand of VgrG17, and the second interaction involved the TTR domain of VgrG17 and a single β-strand of Tae17; this Tae17 β-strand is between the Ig-like domain and the LysM domain. Hydrogen bonding would anchor the length of β-strands from the opposing molecules together, and side chains form further stabilising salt bridges. These salt bridges are mediated by the negatively charged VgrG17 C-terminal region and the positively charged Tae17 N-terminal region. The presence of two interaction sites highlights the importance of a strong interaction for the delivery of the effector, given the mechanical force of T6SS ejection. Further interactions may occur outside these domains, including with the VgrG17 stalk, when the Tae17 effector is loaded for delivery. When we map VgrG17 residues G1069 and W1075 onto the interaction structure, G1069 is found at the end of the TTR domain before a loop region is formed that threads through Tae17. Mutation of this residue to an alanine may disrupt the VgrG17:Tae17 interaction as alanine is more rigid than glycine and may not allow for the loop region to form as readily. In contrast, W1075 is found buried within the Tae17 Ig-like domain, forming hydrophobic interactions that anchor the loop region of VgrG17 to Tae17. Mutation of the tryptophan to an alanine would reduce the size of the residue and disrupt hydrophobic contacts, destabilising the interaction between VgrG17 and the Tae17 Ig-like domain. While it is possible that the loss of these interactions could be due to changes in the folding of VgrG17, we note that the predicted interaction between the VgrG17 TTR domain and Tae17 shows strong structural similarity to that of VgrG and Tle1 from *E. coli* (36). In the *E. coli* VgrG:Tle1 interactions, one side of a β-strand-rich sandwich comprising three anti-parallel β-strands interacts with a small β-strand of the effector, providing the necessary interactions for delivery. Given that Tae17 and Tle1 are different types of cargo effectors but appear to maintain the same interactions with VgrG for delivery, this suggests that this may be a strongly conserved mechanism of cargo effector tethering to the T6SS.

In this study we have shown that Tae17 is a highly modular protein with four domains separated by flexible linkers. This T6SS effector is bifunctional as it possesses both lytic transglycosylase and amidase activity. Delivery of Tae17 is mediated by specific interactions between the Tae17 region encompassing the Ig-like and LysM domains and the C-terminal region of VgrG17. We have precisely defined two VgrG17 amino acids, VgrG17^G1069^ and VgrG17^W1075^, that are crucial for interaction with Tae17 and predict that these allow VgrG17 to thread through the Tae17 structure and allow for the correct alignment of the Ig-like and TTR domains. The AlphaFold2 and PISA interaction model predicted that these residues are located between two edge-to-edge contact interactions with numerous hydrogen bonds and are further stabilized by salt bridges, which together likely anchor the two molecules with sufficient strength to overcome the shear forces associated with T6SS-mediated effector delivery through target cell membranes.

## MATERIALS AND METHODS

### Bacterial strains and growth conditions

Bacterial strains used in this study are listed in Table 1. All strains were cultured in 2YT broth or Lysogeny broth (LB) at 37°C with shaking (200 rpm). For growth on solid media 1% w/v agar was added. Where appropriate, media was supplemented with ampicillin (100 μg/mL), tetracycline (10 μg/mL), kanamycin (50 μg/mL) or carbenicillin (100 μg/mL). For induction or repression of the P*ara*BAD promoter in pBAD30 (29), media were supplemented with 0.4% arabinose or 0.4% glucose, respectively.

### DNA manipulations

All plasmids used this study are listed in Table 2. Primers used for amplification of DNA for cloning/mutagenesis and for Sanger sequencing are listed in Table 3. Isolation of genomic and plasmid DNA, PCR amplification using KOD high fidelity polymerase or Taq polymerase (colony PCR), restriction enzyme digestions, ligations, Sanger sequencing, transformation of chemically competent *E. coli* and transformation of electrocompetent *A. baumannii* via electroporation were performed as described previously (6, 44). For all cloning reactions, ligated products were used to transform *E. coli* chemically competent cells. Transformants containing plasmids with the correct sequence were identified by colony PCR and the sequence of isolated plasmids was confirmed using Sanger sequencing. For some experiments, the constructed plasmid was used to transform *A. baumannii*. To generate recombinant plasmids encoding truncated VgrG17 proteins, VgrG171-1085 (pAL1565) and VgrG171-1074 (pAL1620, Table 2), high fidelity PCR products were generated using primer pairs BAP8874/BAP8842 and BAP8874/BAP9017, respectively, digested with *Sac*I and *Bsp*EI, then ligated in-frame to similarly digested pAL1446, encoding amino acids 1-546 of VgrG17.

**Table 3.**
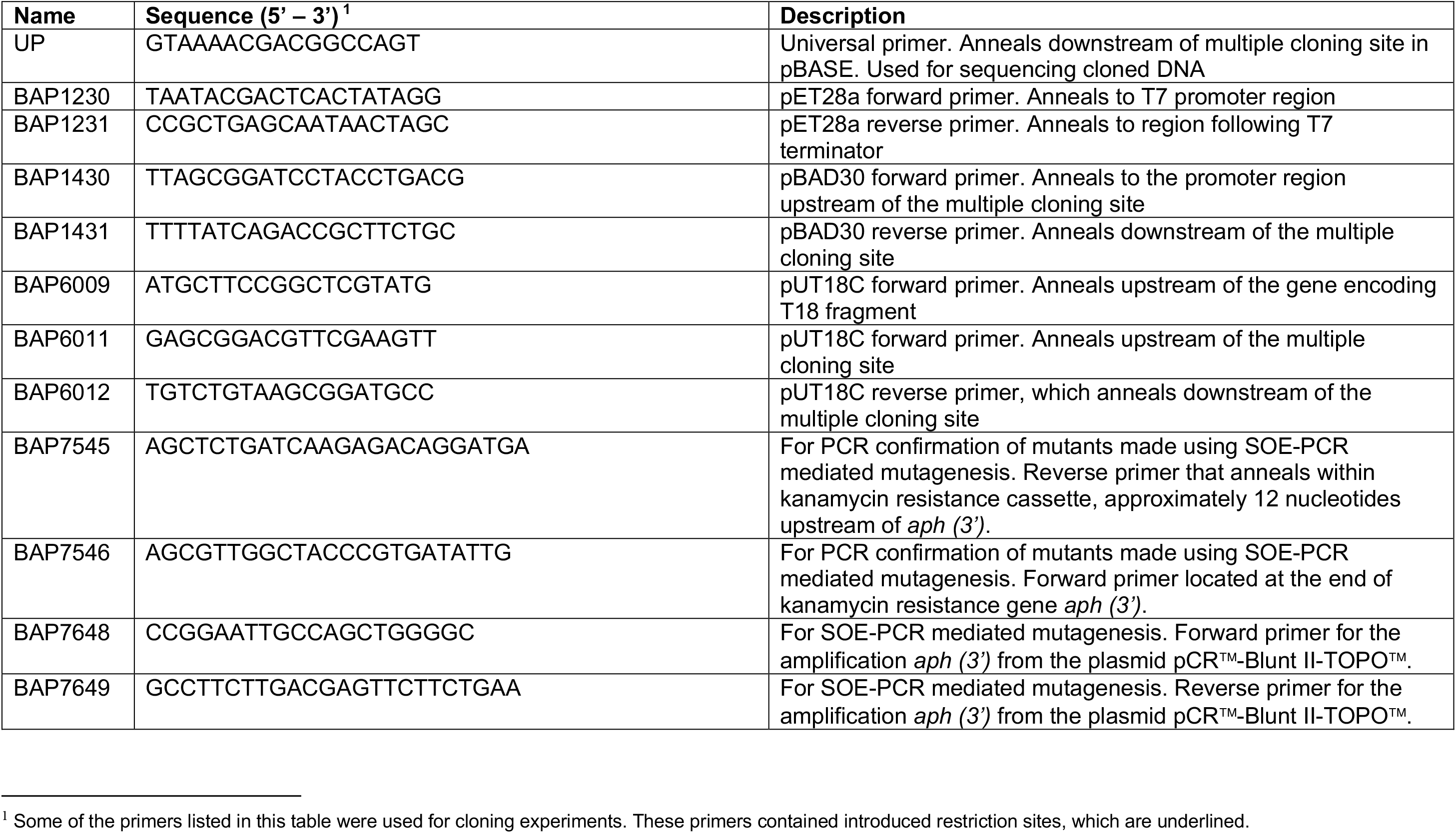
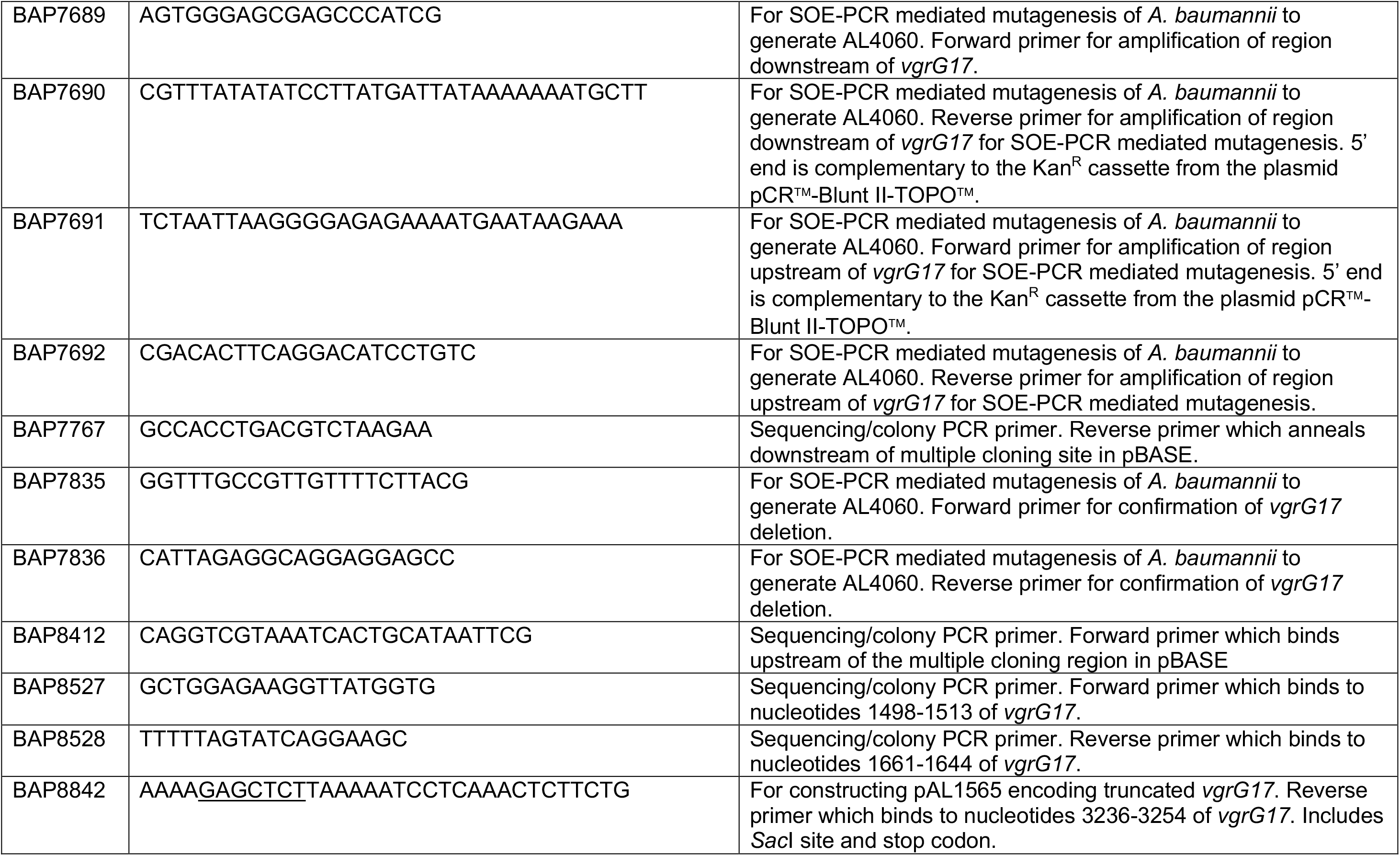
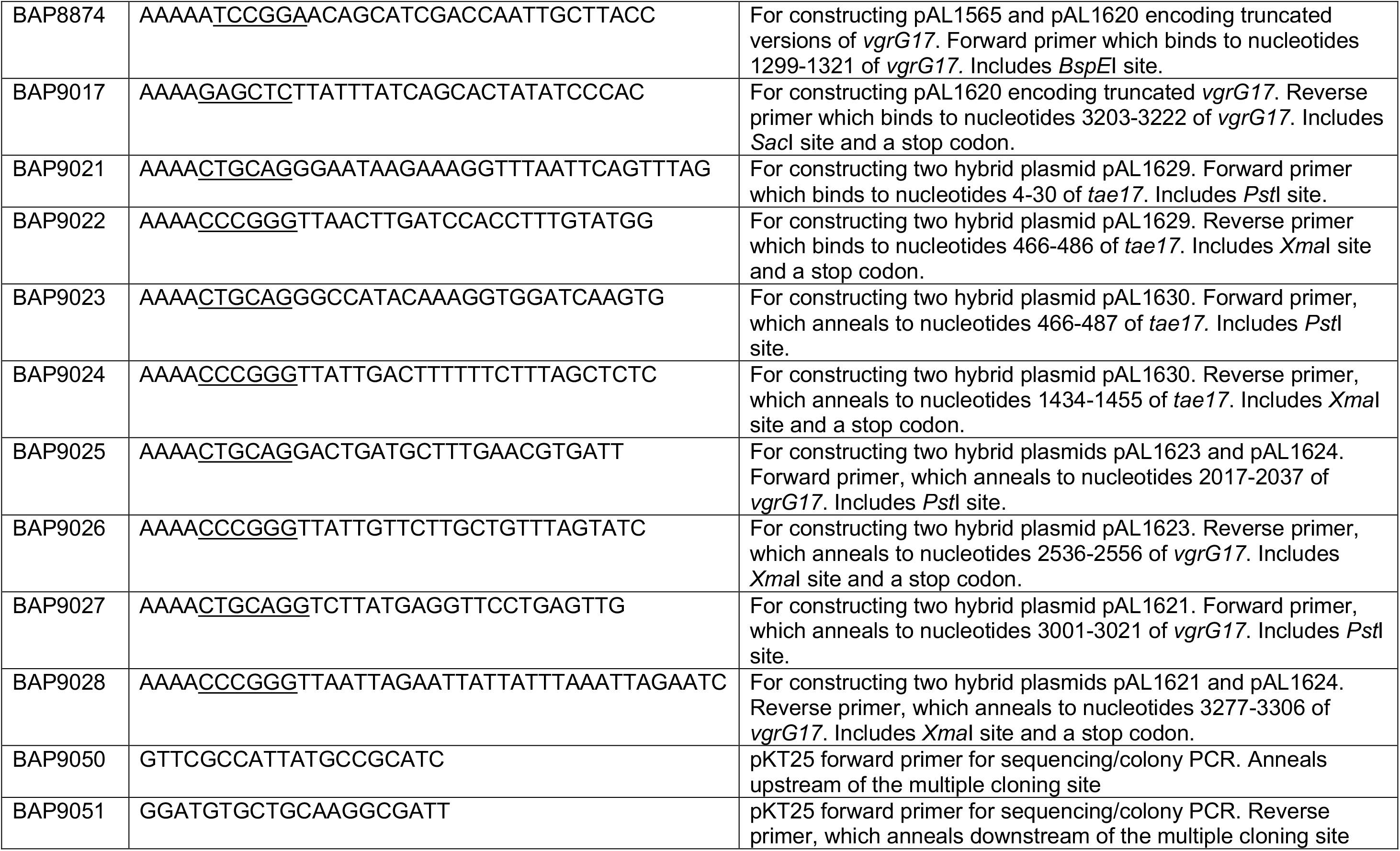
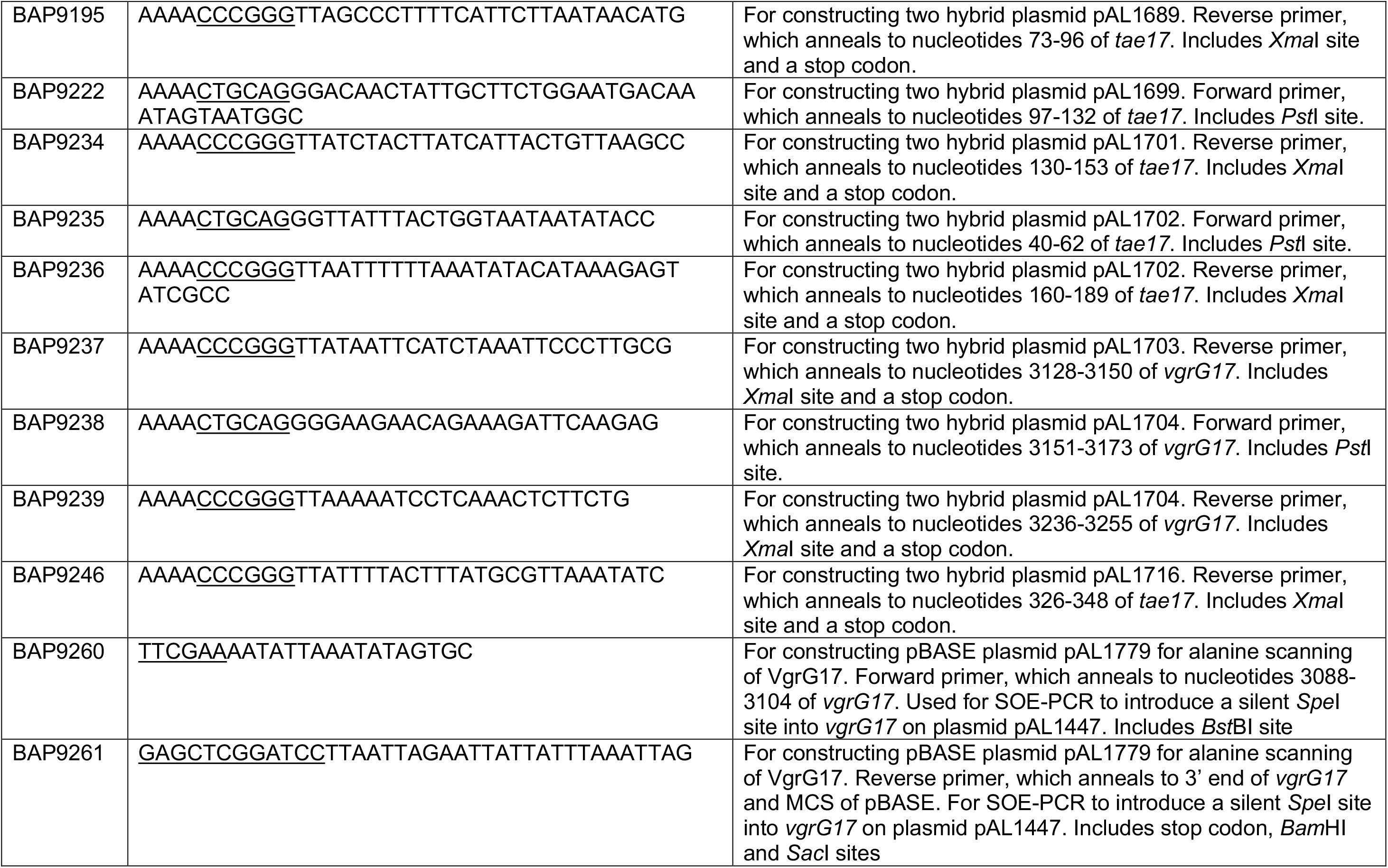
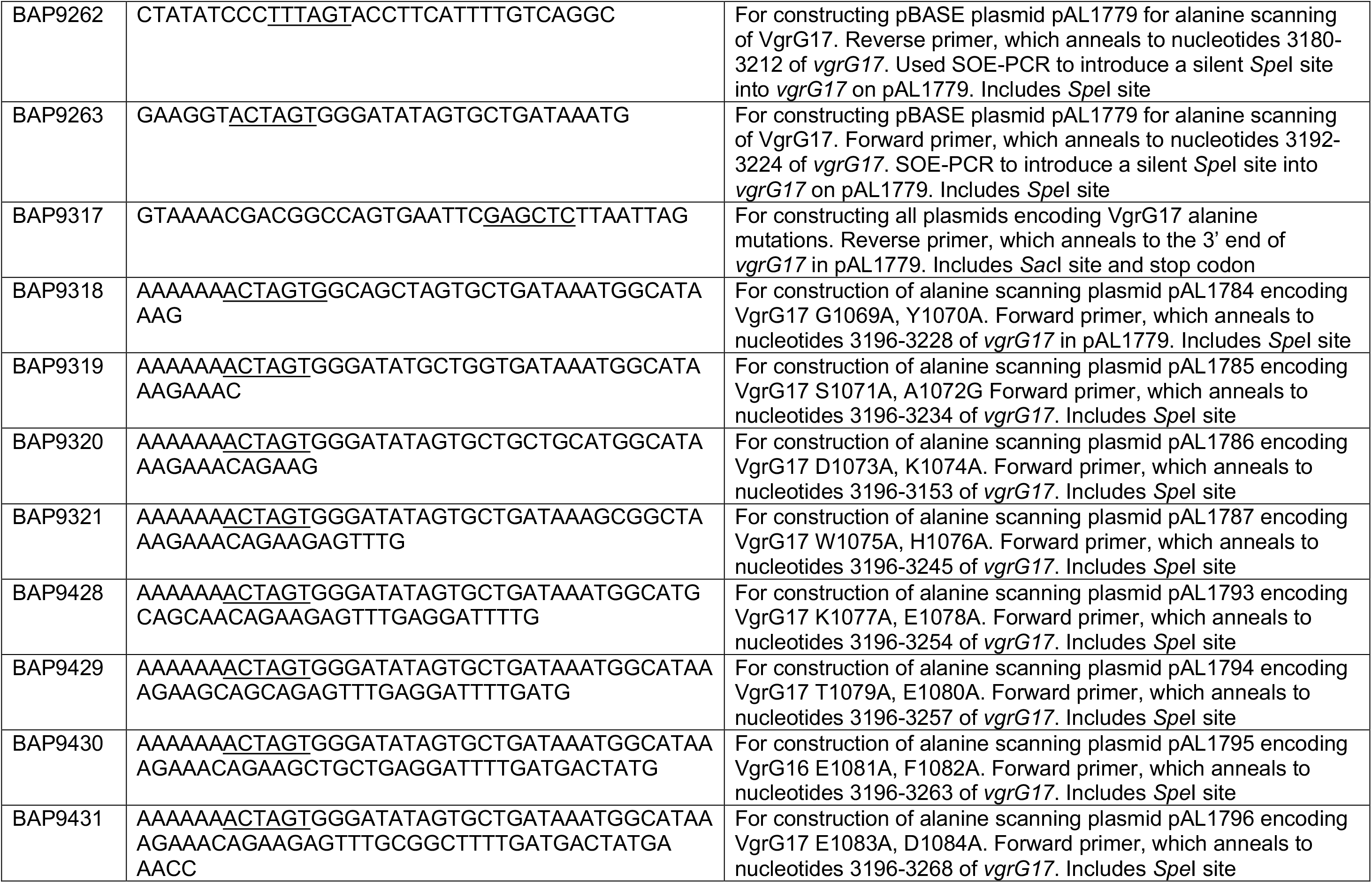
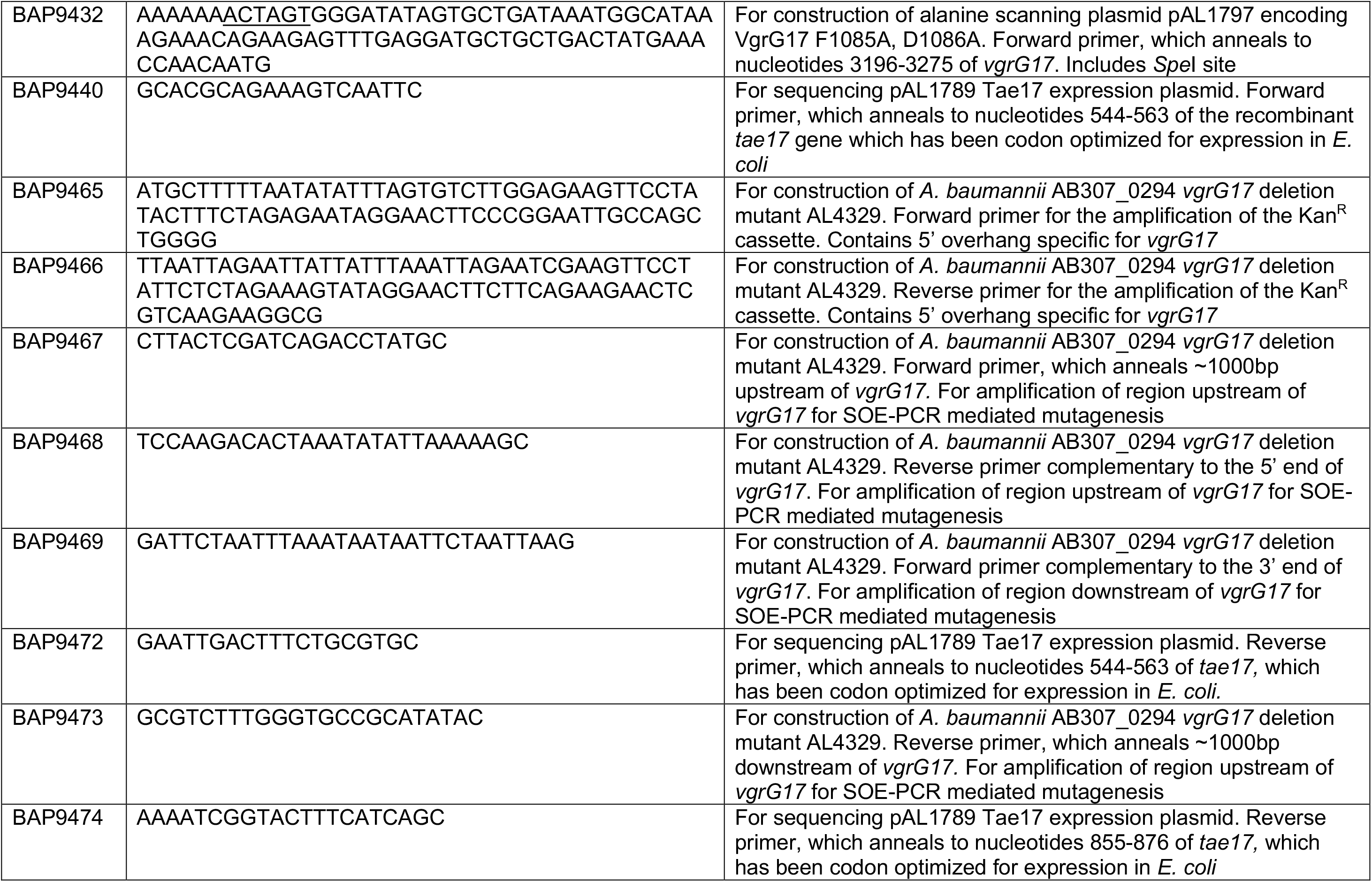
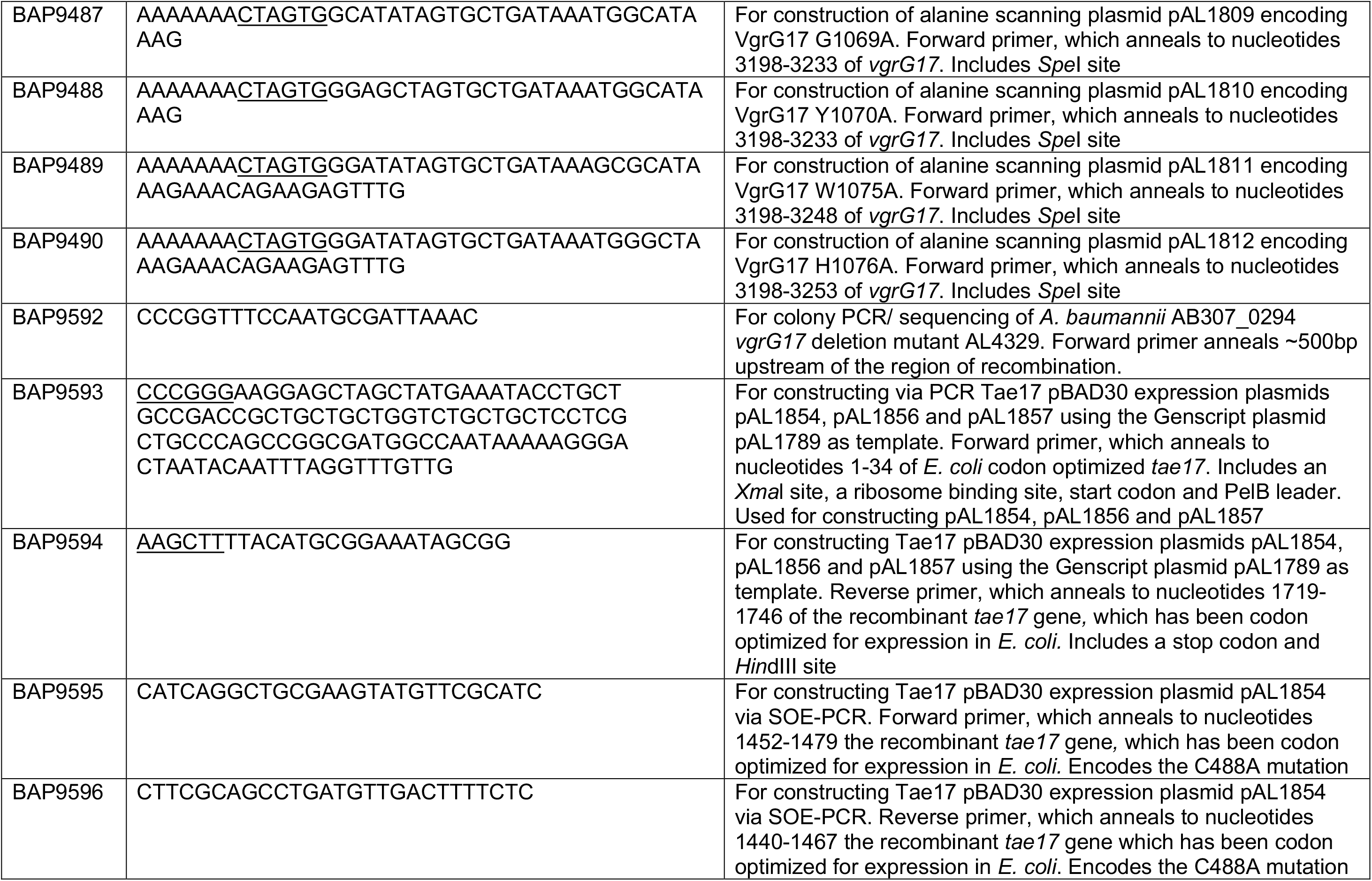
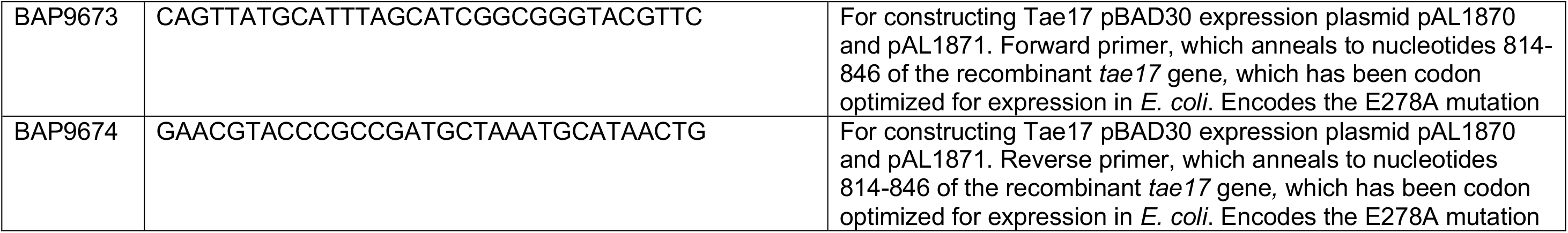
| Primers used in this study.

To construct each plasmid (Table 2) for the adenylate cyclase two-hybrid assay, a region of *vgrG17* or *tae17* was PCR-amplified using a forward primer containing the desired mutations together with the appropriate primer pair, both with suitable restriction sites (Table 3) then ligated to similarly digested pUT18C or pKT25, so that each encoded product was in-frame and at the C-terminal end of the encoded adenylate cyclase fragment T18 or T25, respectively. Pairs of recombinant pUT18C and pKT25 plasmids with the correct in-frame *vgrG17* and *tae17* sequence, respectively, were then introduced into chemically competent *E. coli* BTH101 cells and co-transformants recovered on ampicillin and kanamycin.

For single-site mutagenesis of *tae17* in pBAD30 (pAL1854, pAL1871, Table 2) splicing by overlap-extension PCR (SOE-PCR) (45), using pAL1789 plasmid template and primers encoding the desired change and restriction enzyme sites (Table 3), was employed to generate each DNA fragment encoding a mutated *tae17* gene. Each SOE-PCR product was then restriction enzyme digested and ligated to similarly digested pBAD30. For generation of *tae17* encoding both the E278A and C488A mutations, QuikChange^TM^ site-directed mutagenesis (Agilent Technologies, Inc., Santa Clara, CA) was performed with primer pair BAP9673/9674 to introduce the mutation encoding E278A into the gene using pAL1854 (encoding Tae17^C488A^) as the template. Following digestion with DpnI to remove template DNA, products were introduced into *E. coli* XL-1 Blue to allow for nick repair.

For alanine scanning mutagenesis of recombinant *vgrG17*, the cloning strategy required modification of pAL1447 to introduce a silent SpeI restriction site at the 3’ end of *vgrG17* as follows. An SOE-PCR product spanning *vgrG17* nts 3082 (BstBI) to SacI (in the vector sequence of pAL1447) was made with first round products generated with pAL1447 template and primer pairs BAP9260/BAP9262 and BAP9263/BAP9261. The product was digested with BstB1 and SacI and cloned into similarly digested pAL1447 (Table 2). To generate SOE-PCR products containing mutations for introducing single and double alanine substitutions into VgrG17 we used one of a series of long forward primers containing the desired mutations and an SpeI site (BAP9318-9321, 9428-9432, Table 3), together with reverse primer BAP9317 containing a SacI site (located downstream of *vgrG17* in the vector). Each SOE-PCR product, representing nucleotides 3196-3306 of *vgrG17* with the desired mutations, was then digested and ligated into similarly digested pAL1779.

#### Construction of *A. baumannii* site-directed mutants

*A. baumannii* strain AB307_0294 mutants AL4060 and AL4329 (Table 2) were constructed by allelic replacement of each gene using SOE-PCR products generated with the appropriate primers (Table 3) as previously described (6) with minor modifications as follows. The central kanamycin resistance gene *aph (3’)* was flanked by FRT sites to allow for later excision of the kanamycin cassette leaving an in-frame FRT scar. The SOE product was column-purified and used to transform *A. baumannii* as previously described (46). The kanamycin resistance cassette was removed from the mutation site via the introduction of the plasmid pAT03 (Table 2), which encodes the Flp recombinase (47). The second and third site-directed mutations in AL4060 were constructed following the same procedure as above using SOE-PCR products with the exception that the kanamycin cassette was retained following *vgrG17* deletion in AL4060. All regions of the genome that underwent mutagenesis were subjected to diagnostic PCR using sets of primers flanking the deleted region, followed by DNA Sanger sequencing across the site of mutagenesis.

### His-tagged Tae17 expression and purification

For *in vitro* experiments involving FITC-labelled peptidoglycan and *in vitro* synthesized dansylated lipid II polymers, recombinant Tae17 and mutated Tae17 proteins were produced as follows. Protein expression plasmids (pET28a+), each with an *E. coli* codon-optimized gene encoding C-terminal Hisx6-tagged Tae17^WT^ (pAL1789) Tae17^E278A^ (pAL1919), Tae17^C488A^ (pAL1920) or Tae17^E278A,C488A^ (pAL1885) were constructed by Genscript. *E. coli* C41 cells, each containing a Tae17 expression plasmid, were separately grown in 400 mL of 2YT supplemented with kanamycin at 37°C with shaking at 200 rpm to an OD600nm of 0.6. Recombinant protein expression was induced via the addition of 1 mM IPTG and further incubation at 18°C with shaking (200 rpm) for ∼18 h. Following centrifugation (5,000 *x g*, 20 min) cells were resuspended in 40 mL of a phosphate buffer consisting of 436 mM NaCl, 7 mM K2HPO4, 2.5 mM KH2 PO4 (pH8.0), 5% w/v glycerol (used in all subsequent steps), 20 mM imidazole and Pierce^TM^ Protease Inhibitor (1 mini-tablet per 10ml buffer). Cells were sonicated on ice (Soniprep 150 Plus, 10 mAmp, 6 x 30 sec) then centrifuged (15,000 *x g*, 20 min) to remove insoluble material. His-tagged proteins were purified from each cell lysate by gently mixing lysate with 1ml HisPur^TM^ Ni-NTA resin (Thermoscientific), pre-equilibrated in the above buffer. After loading onto a column for immobilized metal-affinity chromatography (IMAC), the resin was washed several times in the above buffer, then washed several times with phosphate buffer containing 50 mM imidazole. Proteins were eluted with phosphate buffer containing 250 mM imidazole. Elution fractions (1 mL) were concentrated using an Amicon Ultra-15 Centrifugal Filter Unit (Merck) and further purified by size exclusion chromatography using HiLoad 16/600 Superdex 75 column (Cytiva) pre-equilibrated in PBS containing 300 mM NaCl. Fractions with recombinant protein were pooled and concentrated and purity assessed using SDS-PAGE and coomassie blue staining.

### *E. coli* viability assays

Cultures of *E. coli* DH5α containing pBAD30 plasmids encoding Tae17^WT^, Tae17^E278A^, Tae17^C488A^ or Tae17^E278A,C488A^ (all with PelB leader sequences) were grown in LB, supplemented with ampicillin and 0.4% glucose (to supress recombinant protein expression), at 37°C with shaking (200 rpm) to an optical density (OD600nm) of 0.2. Cells were centrifuged (5000 x *g* for 5 min) then resuspended in 30 mL phosphate buffered saline (PBS). Cells were washed once more as above, resuspended in 30 mL of fresh LB supplemented with ampicillin, then divided equally (15 mL) into two sterile flasks; one supplemented with 0.4% glucose and the other with 0.4% arabinose to induce P*ara*BAD promoter and recombinant protein expression. Induced and uninduced cultures were incubated at 37°C as above and OD600nm measurements taken at 0 h, 1 h, 3 h, and 5 h as well as samples to determine CFU (10-fold dilutions, 10 μL aliquots of each dilution spotted onto LB agar supplemented with 0.4% glucose and ampicillin).

### T6SS Interbacterial killing assays

To assess the efficiency of Tae17 delivery by the T6SS, interbacterial killing assays were used. For each assay, an appropriate *A. baumannii* predator strain was used in co-culture on solid media with an *E. coli* DH10B prey strain, harbouring either vector only (pWH1266) or pAL1263 (pWH1266 encoding the immunity proteins Tsi15 and Tdi16 to neutralize the activity of the two other T6SS antibacterial effectors). Assays were performed (≥ 3 biological replicates) as described previously (6), with minor modifications; co-cultures (10 μL) were spotted onto LB agar plates and incubated for 3 h at 37°C. For qualitative interbacterial killing assays (survival or non-survival of *E. coli* following co-culture), each co-culture growth was scraped off the plate and then all collected material added as a primary streak only onto LB agar supplemented with tetracycline to select for surviving *E. coli* prey, and LB agar supplemented with kanamycin (50 μg/mL) or carbenicillin (100 μg/mL) to select for the presence of the *A. baumannii* AB307-0294 predator strain. For quantitative T6SS interbacterial killing assay, each co-culture spot was excised from the agar, resuspended in 1 mL of PBS by vigorous vortexing and then serially diluted in PBS. Aliquots (10 μL) of each dilution were spotted onto LB agar supplemented with appropriate antibiotics (as above) to select for either predator or prey.

### Bacterial two-hybrid adenylate cyclase assays

To identify interacting regions between VgrG17 and Tae17 we used bacterial adenylate cyclase two-hybrid assays as described previously (43, 48).

### Detection of Hcp to assess T6SS activity

For the detection of the T6SS protein Hcp in *A. baumannii* whole-cell lysates and culture supernatants by western immunoblotting, samples were prepared as follows. One mL of mid-log phase *A. baumannii* (OD600nm∼2.0) was centrifuged (5,000 x *g*, 5 min), supernatant collected, and cell pellet resuspended in 100 μL SDS-PAGE sample buffer. The supernatant was concentrated 10x using the Amicon Ultra Centrifugal Filter Units (Millipore), and an equal volume of 2x SDS-PAGE sample buffer added. For loading, samples were heated to 99°C (10 min) then centrifuged (11,000 x *g* for 5 min) to remove insoluble material. A 10 μL aliquot of each sample was subjected to electrophoresis (15% PAGE gel, 100 V for 50 min) and then transferred by western blot to a 0.2 μm nitrocellulose membrane (BioRad) (49). Hcp was detected using a polyclonal chicken anti-Hcp serum (1:500) (6), followed by a secondary HRP-conjugated donkey anti-chicken antibody (1:1000) (Merck Millipore). The western blots were analysed with Clarity Western ECL substrate (BioRad) and visualized on an Amersham Imager 680.

### Peptidoglycan degradation assays

DAP-containing (Gram-negative) peptidoglycan was purchased from Merck and labelled with fluorescein isothiocyanate (FITC) (50). Peptidoglycan degradation assays followed the method from Gurnani Serrano *et al.,* 2021 (51). FITC-labelled peptidoglycan was mixed with 1 µM of protein and buffer A (1 x PBS pH 8.0, 5% glycerol, 300 mM NaCl, 20 mM imidazole) was added with the protein up to a total volume of 100 µL with 5 µL of FITC-labelled peptidoglycan (10 mg/mL). The protein-peptidoglycan mix was incubated at 37°C for 2 h to allow for breakdown to occur. Following this, the samples were added to a filter plate (0.2 µM filter) to quench the reaction and the flow through analysed in a plate reader (Ex. 495 nm and Em. 519 nm). The plate reader then measured the fluorescence against the positive control (Lysozyme) to adjust the gain and a single timepoint measurement was recorded.

### Dansylated lipid II transglycosylase activity assay

Dansylated lipid II (LII) was polymerized with the highly processive monofunctional glycosyltransferase enzyme (MGT) from *S. aureus*. Polymerization occurred by incubating 10 μM of dansylated LII with 0.5 μM MGT for 2 h at 37°C in a reaction mixture consisting of Reaction Buffer (100 mM NaCl, 50 mM HEPES, 10 mM MgCl2), 20% (v/v) dimethylsulfoxide (DMSO) and 600 μM lauryldimethylamine oxide (LDAO). This allowed for polymerization of the LII into a high molecular weight polymer. The reaction was stopped by denaturation of the enzyme at 80°C for 10 min. Tae17^WT^, Tae17^E278A^, Tae17^C488A^ or Tae17^E278A,C488A^ were added separately to the reaction mixture at a concentration of 1 μM (final volume 15 μL) and the reactions were incubated at 37°C for a further 2 h. If the Tae17 species was active, this would allow for the high molecular weight polymer to be broken down. A 3 μL aliquot of 5 x loading dye (10% SDS, 250 mM Tris-HCl pH 6.8, 5% β-mercaptoethanol, 0.02% bromophenol blue) was added to stop the reaction. The samples were loaded onto a Novex^TM^ 16% Tricine gel (Invitrogen^TM^) with anode buffer (100 mM Tris-HCl, pH 8.8) in the main body of the gel tank and cathode buffer (100 mM Tris-HCl, 100 mM Tricine, 0.1 % (w/v) SDS) in the gel cavity. The gel was electrophoresed at 100 V for 2 h and imaged with BioRad ChemiDoc^TM^ MP imaging system using the ethidium bromide setting with a 30 sec exposure.

### Structural modelling and analysis

For *de novo* protein structure models, AlphaFold version 2.1.1 was used on the MASSIVE M3 computing cluster (52). For the Tae17 model, the full-length protein sequence was used as input and monomer mode was used. For the VgrG17:Tae17 interaction model, residues 1010-1101 of VgrG17 and full-length Tae17 were used as sequence input, and multimer mode was used. For both runs the full database was utilized, and the directory set as /mnt/reference/alphafold/alphafold_20211129. Five models were produced, with an unrelaxed and relaxed output for each. For the monomer mode, the relaxed models were ranked on their predicted local distance difference test (pLDDT) scores, and the best pLDDT score was used for model building. The top score for Tae17 full-length modelling was an average pLDDT of 76.5. For the multimer mode, relaxed models were ranked using a predicted template modelling (ipTM) score. The top-ranked model (by ipTM score) was used for analysis. The AlphaFold2-generated Tae17 coordinates are now publicly available on the AlphaFold database under accession A0A5K6CL85 (52). All structures were visualized using UCSF Chimera (v1.15) or ChimeraX (v1.6.1). PDBePISA, an online tool for examining protein interfaces, was used to further assess the interactions between Tae17 and VgrG17 (37).

### Statistical analyses

For *E. coli* survival in interbacterial killing assays, *E. coli* recombinant strain viability counts, and optical density of cultures, statistical significance between samples was determined by log10 transforming the data and performing ordinary one-way ANOVA with Tukey’s multiple comparisons test (GraphPad Prism) with significance defined as *p* < 0.05. For the growth curve experiment, involving recombinant *E. coli* strains, statistical analysis was only performed on the final (5 h) timepoint samples. For the peptidoglycan degradation assays ordinary one-way ANOVA was used with Tukey’s multiple comparisons test (GraphPad Prism) with significance defined as *p* < 0.05.

**Supplemental Material** Supplemental Figures S1-S4 Supplemental Table 1 Supplemental Movie M1

## Acknowledgements

This work was supported by NHMRC grants 1165036 (to JDB and MH) and 1128981 (JDB, SM and MH). VB and BKH were supported by RTP scholarships from Australian Department of Education. This work was supported by the MASSIVE HPC facility (www.massive.org.au) (56). We thank David Steer and the Monash Proteomics and Metabolomics Platform (MPMP) for Mass spectrometry protein identification. We thank Jordan Thompson for construction and sharing of the *A. baumannii* Δ*tssM* mutant strain.

## Data accessibility statement

None

